# Extracellular signalling regulates gastrin transcription through site-specific phosphorylation and nuclear redistribution of Menin

**DOI:** 10.64898/2026.04.07.717082

**Authors:** Uloma B. Elvis-Offiah, Ziyu Wen, Xianxin Hua, Juanita L. Merchant

## Abstract

*The multiple endocrine neoplasia type 1 (MEN1)* gene encodes Menin, a nuclear scaffold protein and tumor suppressor that regulates transcription. It is frequently mutated in endocrine neoplasia. MEN1-gastrinomas are aggressive neuroendocrine tumors (NETs) that arise predominantly in the submucosal Brunner’s glands of the duodenum, an organelle rich in extracellular growth factors. Many duodenal NETs retain wild-type *MEN1* allele and nuclear Menin, suggesting post-translational inactivation of its tumor-suppressor function. The Menin C-terminal domain (CTD) contains a conserved phosphorylation site at Ser487 within the first of three nuclear localization signals (NLS1-3). We hypothesized that extracellular signaling regulates Menin by phosphorylating the CTD at Ser487 blocking its nuclear localization. Using CTD deletion mapping, site-directed mutagenesis, and kinase activation in gastric cell lines, we show that loss of NLS1-3 reduces Menin’s nuclear localization, stability, and repression of *GASTRIN*. Cell stimulation by epiregulin, forskolin, or phorbol ester induced Menin Ser487 phosphorylation and its nuclear translocation, relieving repression of *GASTRIN*. The phospho-mimetic S487D mutant remained cytoplasmic and phenocopied CTD deletion of NLS1-3 sustaining de-repression of *GASTRIN*. These findings showed that Ser487 phosphorylation restricts nuclear accumulation of Menin and functionally links extracellular signaling to post-translational modification of Menin that ultimately contributes to transcriptional derepression and neuroendocrine tumorigenesis.

**GRAPHICAL ABSTRACT:** 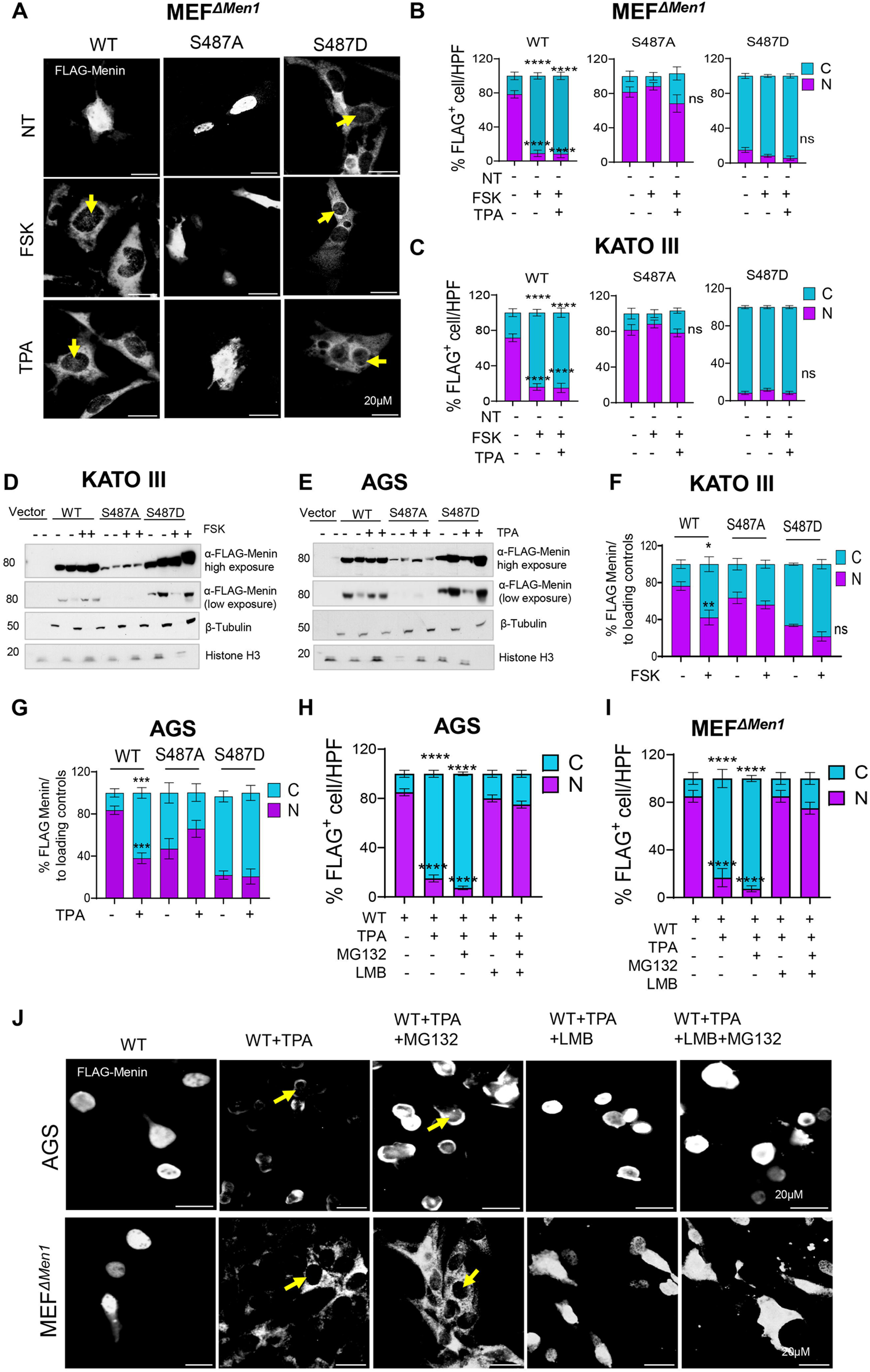

## INTRODUCTION

*Multiple endocrine neoplasia type 1 (MEN1)* is an autosomal dominant cancer syndrome caused by loss-of-function mutations in the *MEN1* gene. It confers susceptibility to endocrine tumors of the pituitary, parathyroid, pancreas, and gastro-duodenal junction, typically manifesting in adulthood [1,2]. *MEN1* encodes Menin, a 610-amino-acid nuclear scaffold protein that interacts with multiple transcription factors and chromatin-modifying complexes to inhibit pro-proliferative genes in neuroendocrine cells, including repression of the *GASTRIN (GAST)* gene [3–6]. Gastrinomas are gastrin-secreting neuroendocrine tumors (NETs) that arise predominantly within the gastrinoma triangle, with approximately 60% originating in the proximal duodenum [2,7]. These tumors represent one of the most aggressive gastroenteropancreatic neuroendocrine neoplasms (GEP-NENs) due to delayed diagnosis, frequent metastasis, and the development of Zollinger–Ellison syndrome (ZES), the physiologic consequence of hypergastrinemia [8–10].

MEN1-associated tumorigenesis has classically been associated with Knudson’s two-hit model, whereby germline inactivation of one *MEN1* allele is followed by somatic loss of the second allele during tumor evolution. However, accumulating clinicopathologic evidence suggests that complete biallelic loss of *MEN1* is not uniformly required for tumor initiation. Notably, up to 50% of small duodenal or pancreatic micro-NETs retain a wild-type *MEN1* allele despite detectable loss of heterozygosity (LOH) at chromosome 11q13 [7]. An MEN1-affected Japanese family harboring a germline exon 10 frameshift mutation (c.1714–1715delTC; p.Ser572Glufs) developed multiple synchronous and metachronous NETs across endocrine organs which retained Menin expression [11]. Immunohistochemical (IHC) analysis of the family members demonstrated progressive loss of nuclear Menin from complete nuclear expression in early lesions to near-absent expression in advanced disease [11]. Collectively, these observations suggest that loss of Menin function may occur in addition to genetic inactivation, implicating additional regulatory mechanism exclusive of LOH.

MEN1-associated duodenal gastrinomas frequently arise from Brunner’s glands (BGs) [12], specialized submucosal structures that a produce growth factor–rich microenvironment of cytokines and mitogens [13,14]. Earlier report shows that BGs express high levels of epidermal growth factor receptor (EGFR) ligands, including amphiregulin (AREG) and Transforming Growth Factor alpha (TGF-α), crucial for protection and regeneration of the intestinal lining [14]. EGFR is a central regulator of epithelial proliferation, survival, migration, and mucosal restitution in the gastrointestinal tract [15]. Its ligand binding induces receptor dimerization and autophosphorylation, activating downstream MAPK/ERK, PI3K-AKT, and JAK/STAT pathways that mediate epithelial repair and crypt cell expansion [16,17]. Moreover, amplification of EGFR family members has been reported in subsets of gastrinomas [18–20]. Consistent with this, we previously demonstrated that EGFR signaling stimulates *GAST* promoter activity and mRNA expression, whereas Menin represses both basal and JunD-mediated transcriptional activation of *GAST* [5,21]. JunD, a component of the AP-1 transcriptional complex, functions as a downstream effector of EGFR-MEK-MAP kinase signaling [22]. These observations suggest that extracellular growth factor signaling may transiently override Menin-mediated repression without *MEN1* gene loss.

The C-terminal domain (CTD) of Menin, encoded by *MEN1* exon 10, contains three nuclear localization signals (NLS1–3) that are essential for its nuclear import [23]. Within Menin’s NLS1 (residues 479–497) is a conserved basic residue–rich motif Arg-Arg-Glu-Ser (RRES) that conforms to a consensus phosphorylation site for cyclic AMP–dependent kinases such as protein kinase A (PKA) [24]. Phosphorylation of Ser487 within this motif has been shown to impair Menin chromatin binding and attenuate repression of *insulin* transcription in pancreatic β-cells [24], but whether this regulatory mechanism operates in neuroendocrine tumors is unknown. Here, we address this gap in understanding by testing the hypothesis that extracellular signals regulate Menin’s cellular location and ultimately function by phosphorylating Ser487. We demonstrate that EGFR activation, cyclic AMP/PKA, and protein kinase C (PKC) signaling pathways phosphorylated Ser487 within NLS, which removes Menin’s repression of gastric gene expression in human gastric cell lines.

## MATERIAL AND METHODS

### Human tissue acquisition

Human tumor specimens, including formalin-fixed paraffin-embedded (FFPE) sections and fresh surgically resected tissues, were obtained in de-identified form from multiple institutional biorepositories, including the University of Arizona Tissue Acquisition Shared Resource (TACMASR), the University of Arizona Tissue Acquisition Repository for Gastric and Hepatic Tissues (TARGHETS), the University of Michigan Endocrine Oncology Repository, the Icahn School of Medicine at Mount Sinai Biorepository and Pathology Core, and the Chao Family Comprehensive Cancer Center Experimental Tissue Shared Resource. Normal human duodenum (nDUO) biopsy specimens were obtained from the University of Arizona Cancer Center (UACC) TARGHETS biorepository. All specimens were collected under protocols approved by the respective Institutional Review Boards (IRB #1808861863, #1909985869, #HUM00115310, #STUDY-12-00145, and #STUDY00000833). Samples were fully de-identified prior to use, and written informed consent was obtained from all participants in accordance with institutional and federal guidelines.

### Antibodies

The following primary antibodies were used for western blotting: anti-FLAG M2 (1:10,000, Sigma-Aldrich, St. Louis, MO, USA, Cat#F3165), phospho-Ser487 Menin (1:200, custom antibody, [24]), PRKX (1:1,000, Invitrogen, Thermo Fisher Scientific, Waltham, MA, USA, Cat#PA5-101865), from Cell Signaling Technology (Danvers, MA, USA) phospho-AKT (Ser473) (1:1,000, Cat#4060), pan-AKT (1:1,000, Cat#11848), CREB (1:1,000, Cat#9197), phospho-CREB (Ser133) (1:1,000, Cat#9198), PKA Cα (1:1,000, Cat#4782), ERK1/2 (1:1,000, Cat#9102), and phospho-ERK1/2 (Thr202/Tyr204) (1:1,000, Cat#9101).

Primary antibodies used for immunohistochemistry (IHC) and immunocytochemistry/immunofluorescence (ICC/IF) included Menin (1:750, Bethyl Laboratories, Montgomery, TX, USA, Cat#A300-105A), and anti-FLAG (1:750, clone D6W5B, Cell Signaling Technology, Cat#14793), EREG (1:200, Santa Cruz Biotechnology, Dallas, TX, USA, Cat#sc-376284), and AREG (1:250, R&D Systems, Minneapolis, MN, USA, Cat#AF262-SP). From Abcam, we purchased EGF (1:300, Abcam, Cambridge, UK, Cat#ab184265), EGFR (1:100, Cat#ab52894), TGFα (1:300, clone EPR15346, Cat#ab208156), synaptophysin (1:1,000, Cat#ab16659).

Secondary antibodies and antibodies for loading controls purchased from Cell Signaling were GAPDH (clone D16H11) (1:1,000, Cat#5174), β-tubulin–HRP (1:5,000, Cat#5346), Histone H3–HRP (1:10,000, Cat#12648), Lamin B1 (1:2,000, Cat#13435), from Thermo Fisher, we purchased donkey anti-rabbit IgG Alexa Fluor 488 (1:400, Cat#A-21206), donkey anti-rabbit IgG Alexa Fluor 594 (1:400, Cat#A-21207), donkey anti-rabbit IgG Alexa Fluor 647 (1:400, Cat#A-31573), donkey anti-mouse IgG Alexa Fluor 488 (1:400, Cat#A-21202), and donkey anti-mouse IgG Alexa Fluor 594 (1:400, Cat#A-21203).

### Plasmid Design and Purification

Plasmid design and construction were based on the human MEN1 gene sequence originally reported by Chandrasekharappa et al. (GenBank accession U93236). The wild-type *MEN1* construct (Clone ID: OHu17935D) corresponds to the NCBI RefSeq transcript NM_130799.2, encoding a full-length 1830 bp open reading frame (ORF). Plasmid construction and sequence validation were performed by GenScript Biotech Corporation (Piscataway, NJ, USA). The *MEN1* coding sequence was cloned into the BamHI and XbaI restriction sites of the pcDNA3.1(+)/C-(K)-DYK expression vector (Thermo Fisher Scientific,), placing the ORF upstream of a C-terminal FLAG tag (DYKDDDDK) to enable detection of the expressed protein. In parallel, additional *MEN1* constructs carrying sequential C-terminal truncations (∼60 bp; 20 amino acids) were generated and included in comparative analyses (see Supplementary Table 1). All constructs were verified by Sanger sequencing. Plasmids were reconstituted in Tris-EDTA (TE) buffer prior to bacterial transformation. Chemically competent DH5α Escherichia coli cells (Thermo Fisher Scientific) were transformed with the pcDNA3.1(+) empty vector, wild-type *MEN1*, or corresponding mutant constructs. Transformants were selected on LB agar plates containing 100 µg/mL ampicillin. A single colony was inoculated into LB broth supplemented with ampicillin and cultured overnight at 37°C with shaking at 250 rpm. Plasmid DNA was purified using the QIAGEN Plasmid Plus Midi Kit (QIAGEN, Hilden, Germany, Cat#12943) according to the manufacturer’s instructions. Purified plasmids were eluted in sterile TE buffer and stored at -20°C until use.

### Site Directed Mutagenesis

In addition, two Ser487 phosphorylation-specific MEN1 mutant plasmids (S487A and S487D) were generated by GenScript Biotech Corporation (see Supplementary Table 1). Mutations were introduced using a high-fidelity PCR-based proprietary mutagenesis strategy, followed by DpnI digestion and clonal selection according to the manufacturer’s standard protocol. All constructs were verified by Sanger sequencing to confirm the intended mutation and to exclude off-target nucleotide changes. Plasmids were transformed into DH5α Escherichia coli cells, amplified, and purified as previously described. DNA was reconstituted in nuclease-free water and validated in-house prior to transfection.

### Cell Culture

Human gastric cancer cell lines expressing unprocessed gastrin peptide, AGS (gastric adenocarcinoma; #CRL-1739) and KATO III (gastric carcinoma; #HTB-103) were obtained from the American Type Culture Collection (ATCC, Manassas, VA, USA), and MKN-45 (gastric carcinoma; #JCRB0254) was obtained from the Japanese Collection of Research Bioresources (JCRB, Osaka, Japan). AGS cells were maintained in Ham’s F-12 medium supplemented with 5% fetal bovine serum (FBS) and 1% penicillin–streptomycin (Corning Inc., Corning, NY). MKN-45G cells were cultured in RPMI-1640 medium containing L-glutamine, supplemented with 5% FBS and 1% penicillin–streptomycin. KATO III cells were maintained in Iscove’s Modified Dulbecco’s Medium (IMDM) supplemented with 10% FBS and 1% penicillin–streptomycin. Cells were mainta ined at 37 °C in a humidified incubator with 5% CO₂. *Men1*-null mouse embryonic fibroblasts (MEF*^ΔMen1^*) were kindly provided by Dr. S. K. Agarwal (NIH, Bethesda, MD, USA). They were cultured in Dulbecco’s Modified Eagle Medium (DMEM) supplemented with 10% fetal bovine serum and penicillin–streptomycin. All cells were incubated at 37°C in a humidified atmosphere with 5% CO₂ and used between passage numbers 3 and 10. Cells were routinely tested for Mycoplasma contamination every six months using the Universal Mycoplasma Detection Kit (ATCC, Cat#30-1012K) before initiating experiments.

### Gastrin-Luciferase Reporter Assay

The gastrin-luciferase (240 GASLuc) reporter system was used to evaluate transcriptional activity at the *GAST* promoter, as previously described [5,25]. A 240 bp sequence from the human *GAST* gene promoter was subcloned upstream of the firefly luciferase coding sequence in the pGL3-Basic Luciferase Reporter Vector (Promega, Madison, WI, USA, #E1751). AGS, KATO III, and MKN45 cells were seeded in 6-well plates. Upon reaching 60%–75% confluency, these cell lines were co-transfected with 0.75 µg of the 240 GASLuc plasmid and the respective FLAG-tagged *MEN1* expression constructs using jetOPTIMUS transfection reagent (Polyplus-transfection, Illkirch, France). To normalize transfection efficiency, cells were also co-transfected with the PRL-TK Renilla Reporter Plasmid (Promega, #E2241). After 24 h, cells were serum-starved for 24 h, followed by treatment with either recombinant human eregulin (EREG) protein (10 nM), a potent ligand for EGFR (R&D systems, #1195-EP) or 10 µM Forskolin (FSK, ThermoFisher, #66575-29-9), activator of adenylyl cyclase and cyclic AMP or and 10nM phorbol 12-myristate 13-acetate (TPA; Sigma-Aldrich, #P8139) for 4-8 h. Cells were lysed, and luciferase activity was measured using the Dual-Luciferase Reporter Assay System (Promega, #E1980) according to the manufacturer’s instructions. Luminescence was quantified using a Synergy 2 Microplate Reader and analyzed with Gen5 software (BioTek instruments, Winooski, VT, USA).

### RNA Extraction and Quantitative RT-PCR

To evaluate *MEN1* and *GAST* mRNA expression, cells were serum-starved for 24 h in 1% FBS and then transfected with either wild-type or mutated Menin. After additional 24 h, cells were treated without or with either EREG (10 nM) or FSK (10 µM) or TPA (10 nM) for 2-4 h before lysis and RNA extraction. Total RNA was extracted using the PureLink RNA Isolation Kit (Invitrogen, #12183025). Cells were lysed in RNA lysis buffer supplemented with 1% β-mercaptoethanol (Sigma-Aldrich, #M6250) and homogenized by passing the lysate through a 20-gauge syringe needle seven times. The lysate was mixed with an equal volume of 70% ethanol and applied to PureLink spin columns for RNA purification according to the manufacturer’s protocol. Up to 1 µg of total RNA was treated with ezDNase enzyme mix at 37°C for 2 min to remove genomic DNA contamination, followed by cDNA synthesis using SuperScript™ IV VILO Master Mix (Thermo Fisher Scientific, #11756050). Quantitative RT-PCR (qRT-PCR) was performed using PowerUp™ SYBR Green Master Mix (Applied Biosystems, #A25742), with 5 ng of cDNA per reaction. Reactions were run on a Bio-Rad CFX thermocycler using the following conditions: 95°C for 2 min, followed by 40 cycles of 95°C for 15 seconds and 60°C for 1 minute. All forward and reverse primers were obtained as validated predesigned PrimeTime qPCR Assay primer sets for SYBR Green chemistry (Integrated DNA Technologies, Coralville, IA, USA) and used at a final concentration of 500 nM per primer. Relative gene expression was calculated using the 2^⁻ddCt^ method and then log transformed [26].

### Western Blot Analysis

Protein extracts (20–25 µg per lane) were prepared under reducing conditions using 1× SDS loading buffer supplemented with 5% β-mercaptoethanol and denatured by boiling at 95 °C for 5 min. Proteins were separated on precast 4–12% Bis-Tris gradient gels (Invitrogen, #NP0322BOX) using 1× MOPS electrophoresis buffer at 90 V for 10 min followed by 110 V for 90 min. Gels were transferred onto polyvinylidene difluoride (PVDF) membranes using the iBlot 2 Dry Transfer System (Invitrogen, #IB21001). Membranes were blocked for 1 h at 24 °C in 5% (w/v) bovine serum albumin (BSA) dissolved in TBS with 0.05% Tween-20 (TBST) and incubated overnight at 4 °C with primary antibodies diluted in 5% BSA-TBST (See antibody section). After three washes in TBST, membranes were incubated for 1 h at 24 °C with HRP-conjugated secondary antibodies followed by three additional TBST washes. Protein bands were detected using the Pierce™ ECL Western Blotting Substrate (Thermo Fisher, #32106) and imaged on autoradiographic film. When required, membranes were stripped and reprobed for loading controls under identical blocking conditions. For quantitation, films were scanned in grayscale and analyzed using FIJI (ImageJ, NIH, Bethesda, MD) software. Bands were identified by expected molecular weight, and equal-area regions of interest (ROIs) were selected. Mean gray values (MGVs) were inverted (255 - MGV), background-subtracted, and normalized to corresponding loading controls. Global brightness adjustments, when applied, were performed uniformly across the entire image.

### Subcellular Fractionation

Within 24 h of seeding cells into 6-well plates, AGS, MKN-45G, and KATO III cell lines were transfected with 1.2 μg of FLAG vectors using jetOPTIMUS. After 24-48 h, cells were washed twice with PBS, detached by mechanical dissociation or after 1 min incubation with Trypsin-EDTA solution (Invitrogen). The cells were collected by centrifugation at 500 × g for 5 min. Pellets were washed once with PBS, and processed for subcellular fractionation using the NE-PER Nuclear and Cytoplasmic Extraction Kit (Thermo Fisher, #78833) according to the manufacturer’s instructions. For cytoplasmic extraction, pellets were resuspended in 0.1 mL Cytoplasmic Extraction Reagent I supplemented with protease/phosphatase inhibitors. The remaining nuclear pellets were extracted with 0.05 mL Nuclear Extraction Reagent to obtain nuclear proteins.

### Immunohistochemistry (IHC)

Human tissue specimens were fixed overnight at 4 °C in 4% paraformaldehyde, processed, and embedded in paraffin. Paraffin blocks were sectioned at 5 µm thickness and mounted onto frosted glass slides. Slides were deparaffinized with three washes in xylene and rehydrated through decreasing ethanol concentrations (100%, 90%, and 70%) followed by rinsing in phosphate-buffered saline (PBS). Sections were washed in TBST (Tris-buffered saline containing 0.1% Triton X-100) prior to heat-mediated antigen retrieval, which was performed using Tris-EDTA buffer (pH 9.0; Abcam, # ab93684) for 30 min, followed by cooling at room temperature for 15 min and subsequent washing in TBST. Endogenous peroxidase and alkaline phosphatase activities were blocked using BLOXALL blocking solution (Vector Laboratories) for 10 min, followed by three washes in TBST. Sections were then blocked for 1 h at room temperature in 10% donkey serum prepared in TBST supplemented with 1% bovine serum albumin (BSA) and 0.2% Triton X-100. Primary antibodies were incubated overnight at 4 °C at the dilutions specified in the Antibodies section. Slides were washed in TBST and incubated with anti-rabbit IgG, horseradish peroxidase (HRP)–conjugated secondary antibody (Cell Signaling, Cat# 7074) diluted 1:300 in TBST containing 1% BSA for 1 h at room temperature. Signal detection was performed using 3,3′-diaminobenzidine (DAB; Vector Laboratories, Burlingame, CA) followed by counterstaining with hematoxylin (Leica Biosystems, Buffalo Grove, IL, USA). Slides were then dehydrated through increasing ethanol concentrations (70%, 90%, and 100%), cleared in xylene, and mounted with a permanent mounting medium. Images were acquired using an Olympus BX53F epifluorescence microscope equipped with an Olympus digital camera (Olympus, Center Valley, PA) at 100×, 200×, and 400× magnifications. Image acquisition settings were maintained consistently across samples for comparative analysis. Post-acquisition image processing was performed using FIJI/ImageJ software, with adjustments applied globally to entire image files and limited to channel-level modifications. Immunoreactivity for TGFα, and EREG was semi-quantitatively assessed using an H-score method, calculated as: H-score = ∑(percentage of cells at each intensity × intensity [0–3]), yielding a range of 0-300. Cytoplasmic staining intensity was evaluated in BGs and tumor cells across multiple representative fields per case. Menin immunoreactivity was assessed by quantifying nuclear and cytoplasmic staining across multiple high-power fields, with a minimum of 300 cells evaluated per case. Nuclear positivity was defined as distinct DAB signal within the nucleus, whereas cytoplasmic positivity was defined as diffuse or granular staining exceeding background. Nuclear-to-cytoplasmic (N/C) ratios were calculated for each sample.

### Immunofluorescence (IF)

MEF*^ΔMen1^*, AGS and KATO III cells were seeded onto glass coverslips and incubated overnight to allow for adherence. Cells were transfected with FLAG-tagged wild-type_*MEN1 and* mutant plasmids and serum-starved as previously described. At 24 h post-transfection, cells were treated with or without EREG (10 nM) or FSK (10 µM) for 2-4 h before fixation. Cells were washed twice with PBS to remove serum and fixed in 4% paraformaldehyde (Electron Microscopy Sciences, #15710) for 20 min at 24 °C. After fixation, cells were washed three times in PBS for 5 min per wash and permeabilized with 0.5% Triton X-100 in TBST for 10 min at 24 °C. Coverslips were transferred onto prelabeled clean slides and blocked for 1 h at 24 °C in blocking buffer containing 10% donkey serum (Jackson ImmunoResearch, #017-000-121), 0.1% BSA, and 0.1% Triton X-100 in TBST. Cells were then incubated overnight at 4 °C in a humidified chamber with 1: 800) rabbit anti-FLAG. Following three washes in TBST, coverslips were incubated with secondary antibody for 1 h at 24 °C in a humidified chamber. Coverslips were washed in TBS and stained with Alexa Fluor® 488-conjugated phalloidin (1:100; Cell Signaling, #8878) or Alexa Fluor® 488-conjugated anti-β-Tubulin (clone 9F3) (1:200; Cell Signaling, #3623) to visualize the cytoplasmic compartment. Following washes in TBS, cells were mounted using ProLong Gold Antifade with DAPI (Invitrogen; #P36931) and imaged using an Olympus epifluorescence microscope equipped with cellSens software. For quantification of nuclear versus cytoplasmic Menin localization, 5 to10 fields at 400× magnification were captured per plasmid condition per experimental replicate, yielding 15 to 30 images per construct. Menin localization was classified based on colocalization with DAPI, with nuclear localization defined by signal overlap with DAPI and cytoplasmic localization defined by exclusion from the nuclear compartment. Margins were defined by using images of wild-type Menin, which displayed crisp, rounded nuclear morphology. Statistical analysis was performed using GraphPad Prism 10 software.

### Kinase prediction analysis

The dataset employed was retrieved from the public PhosphoSitePlus Kinase Library database (https://www.phosphosite.org/kinaseLibraryAction.action?siteId=18012115, accessed on 7 June 2023) and comprises a curated collection of human protein kinases annotated with motif-based compatibility scores for the Menin Ser487 phosphorylation site. The amino acid sequence spanning the NLS1 region surrounding Ser487 was queried using default stringency parameters to identify candidate serine/threonine kinases. Kinases were ranked by Site Percentile, Percentile Rank, and activity score, with higher percentile values indicating stronger predicted motif compatibility. Prediction results were used to support hypothesis generation and to contextualize experimental signaling data and were not interpreted as definitive evidence of direct kinase–substrate relationships.

### Statistical Analysis

All statistical analyses were performed using GraphPad Prism 10. For comparisons between two groups, unpaired Student’s t-tests were applied. One-way ANOVA with Tukey’s post hoc test was used for comparisons among three or more groups. Two-way ANOVA with Tukey’s post hoc test was used when evaluating multiple groups across more than one condition. For time-course assays, pairwise comparisons across plasmid conditions were restricted to each respective timepoint. Quantitative PCR and luciferase assays were conducted in three or more independent biological replicates, with data normalized to the negative control for each experiment to account for technical variation. Data are presented as mean ± SEM.

## RESULTS

### *GAST* repression requires intact Menin NLSs

Despite the prediction that Menin contains three NLSs to mediate nuclear import, evidence that they modulate Menin’s location has not been clearly established. For example, in mouse embryo fibroblasts, NLS point mutations did not modulate Menin’s cellular location despite impairing insulin binding protein gene expression [23,27]. Others have reported that a truncated CTD decreases Menin stability and renders it non-functional by increasing its aggregation and degradation [28]. Although we and others have demonstrated that Menin can shuttle in and out of the nucleus [29,30], dependence on the NLSs was not directly assessed. Nevertheless, patients harboring C-terminal truncated forms of Menin encoded by exon 10, exhibit severe NET phenotypes and cytoplasmic expression [11,28,31]. Therefore to define whether these NLS motifs regulate Menin’s location and repressive function on *GAST* gene expression, we performed scanning mutagenesis of Menin’s CTD by deleting the region spanning 470 to 610 in 20–amino acid increments (**Fig. 1A**). To assess the transcriptional repressor activity of the to sequentially deleted CTDs, we measured *GAST* gene expression using a previously characterized human luciferase (Luc) reporter expressed from the first 240 bp of the human *GAST* promoter (240*GASLuc*) [5,32]. Gastrin-expressing human gastric cell lines (KATO III, AGS, and MKN-45G) were co-transfected with wild-type Menin or the CTD deletion constructs, the 240 *GASLuc* reporter and a *Renilla* Luc control vector. Under basal conditions, wild-type Menin and all mutants retaining at least one intact NLS significantly suppressed *GAST* promoter-driven Luc activity by ∼50–60% relative to the empty vector (**Fig. 1B**). In contrast, mutants lacking all three NLSs showed a reduction in *GAST* reporter activity (70-90%) relative to wild-type Menin, with the Luc activity approaching levels observed in the absence of exogenous Menin, indicating loss of CTD-dependent repressive function (**Fig. 1B**). Since these gastric cell lines also express endogenous *GAST*, quantification of endogenous *GAST* mRNA by qRT-PCR following transient transfection demonstrated a similar loss of Menin repression by the constructs lacking NLS1–3 (**Fig. 1C**). Thus, both the *GASLuc* reporter activity, and endogenous *GAST* mRNA results confirmed that all three Menin NLSs are required for transcriptional repression.

**Figure 1.**
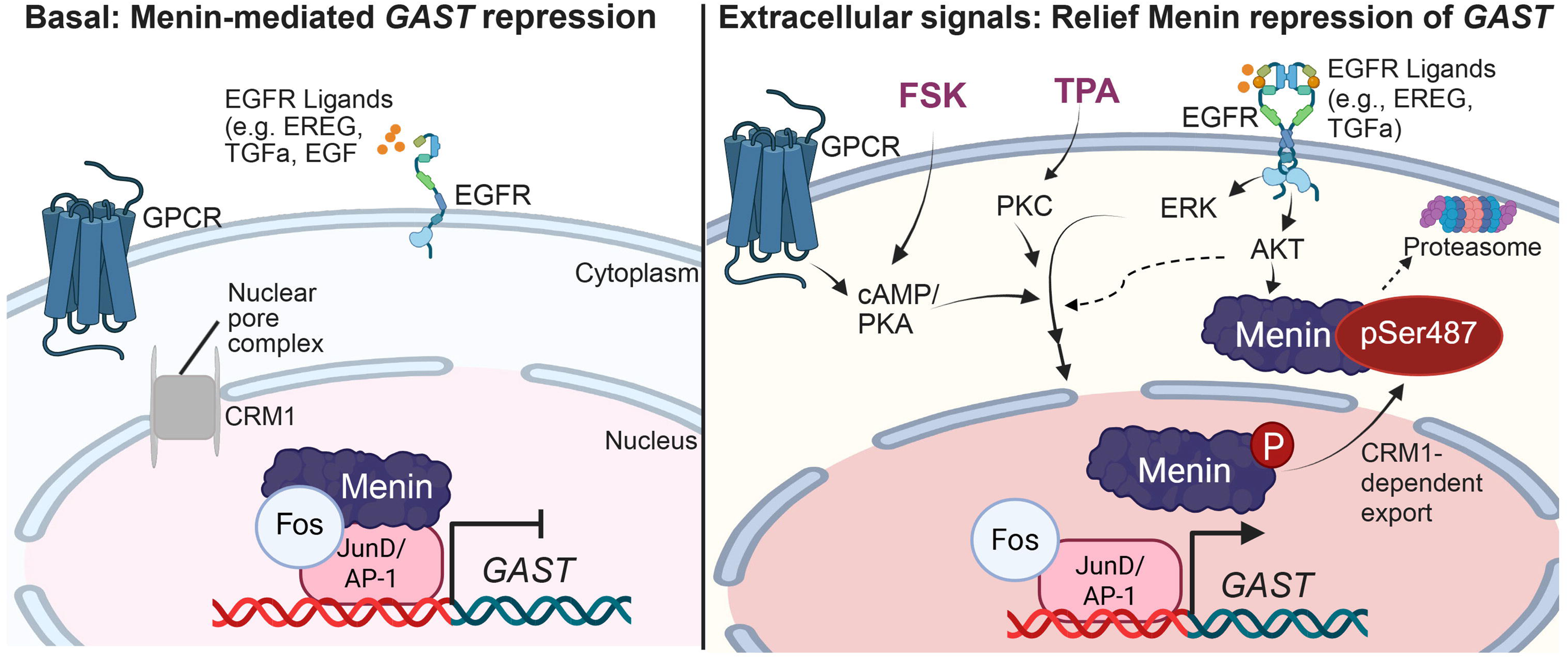
Loss of the Menin CTD impairs *GAST* repression in gastric carcinoma cells. (A) Schematic of FLAG-tagged C-terminal Menin deletion constructs from amino acids 470 to 610. (B) Luciferase (Luc) reporter activity in AGS, KATO III, and MKN-45G cells co-transfected with the 240 GasLuc and either empty vector, wild-type Menin, or CTD deletion constructs, normalized to *Renilla* Luc activity. (C) Relative mRNA expression of endogenous *GAST* in AGS, KATO III, and MKN-45G cells following transfection with empty vector, wild-type Menin, or deletion constructs, normalized to *HPRT1*. (D) Relative *MEN1* mRNA expression in AGS, KATO III, and MKN-45G cells following transfection with empty vector, wild-type Menin, or deletion constructs, normalized to *GAPDH*. (E) Representative Western blot analysis of total Menin protein levels in AGS, KATO III, and MKN-45G cells under basal conditions; β-tubulin served as a loading control. (F) Quantification of FLAG–Menin protein expression normalized to β-tubulin. Data are presented as mean ± SEM from three independent biological replicates. Luciferase and qRT-PCR data represent n = 3–5 independent expts, performed in triplicate. Significance was determined by one-way ANOVA with Tukey’s post hoc test; *P < 0.05, **P < 0.01, ***P < 0.001, ****P < 0.0001; ns, not significant.

To determine whether the observed changes in *GAST* gene expression were due to altered Menin protein expression, we quantified *MEN1* mRNA and Menin protein levels between wild-type and CTD deletion constructs. Quantitative RT-PCR revealed that *MEN1* transcript levels were comparable across all constructs and cell lines (**Fig. 1D**). In MKN-45G cells, *MEN1* mRNA expressed from the cytomegalovirus (CMV) promoter (pcDNA vector) was approximately 70-fold higher than empty vector, whereas AGS and KATO III cells showed a more modest ∼7-fold higher expression; importantly, transcript levels did not significantly differ between wild-type and deletion mutants under basal conditions (**Fig. 1D**). However, Menin protein stability varied among constructs. The C490 and C470 mutants, which disrupt NLS1-3, showed reduced protein expression within 24–48 h post-transfection compared to wild-type and the constructs retaining NLS2 and NLS3, and reached statistical significance in KATO III cells (**Fig. 1E, and F**). To determine whether this decrease reflected proteasome-mediated degradation, cells were pretreated with the proteasome inhibitor 10 nM MG132 for 1 h, a dose previously shown to effectively block proteasomal turnover in gastric cell lines [25,30]. MG132 pretreatment had minimal effects on Menin protein levels across most constructs, whereas the C490 and C470 deletion mutants increased their stability. Collectively, these findings suggest that the reduced suppression from NLS-deficient Menin arises from proteasome degradation of truncated mutants rather than reduced *MEN1* expression.

### Ligand-enriched EGFR signaling in DNETs correlates with cytoplasmic redistribution of Menin

Since Menin-dependent repression of *GAST* requires nuclear localization, we hypothesized that extracellular signaling within the normal duodenal niche may transiently inactivate Menin by promoting its translocation to the cytoplasm in vivo. We therefore demonstrated the localization of two EGFR ligands and Menin in normal human duodenum, Brunner’s glands (BG) and DNETs. EGFR is activated by multiple ligands, including EGF, TGFα, HB-EGF, AREG, BTC, EREG, and epigen [33]. Among these, TGFα is expressed in approximately 70-100% of neuroendocrine neoplasms, with increased expression observed in larger and higher-grade lesions [34,35]. The EGFR is co-expressed with TGFα in up to 80% of cases, supporting a ligand-driven autocrine signaling axis [35,36].

We used hematoxylin and eosin (H&E) staining to define the mucosal architecture and anatomical context for ligand localization, including intestinal crypts and submucosal BGs **(Fig. 2A**). Synaptophysin (SYP), a marker of NET, confirmed the presence of submucosal DNETs (**Fig. 2B**). We observed that in both normal and tumor-adjacent BGs, TGFα expression was reduced and primarily cytoplasmic with weak membranous staining (**Fig. 2C, E**). By contrast, the tumors exhibited strong TGFα expression in a patchy distribution across tumor cell clusters (**Fig. 2C, E**). A similar spatial pattern was also observed for EREG across epithelial, BG, and tumor compartments (**Fig. 2D, F**). In contrast, EGFR expression in normal duodenum was restricted to the villi and primarily membranous, with minimal expression in BGs and stroma. As expected [37], EGFR immunoreactivity in tumors was strong and predominantly cytoplasmic and vesicular, with only focal membranous staining, consistent with receptor internalization and ligand-driven signaling (**Fig. 2G**).

**Figure 2.**
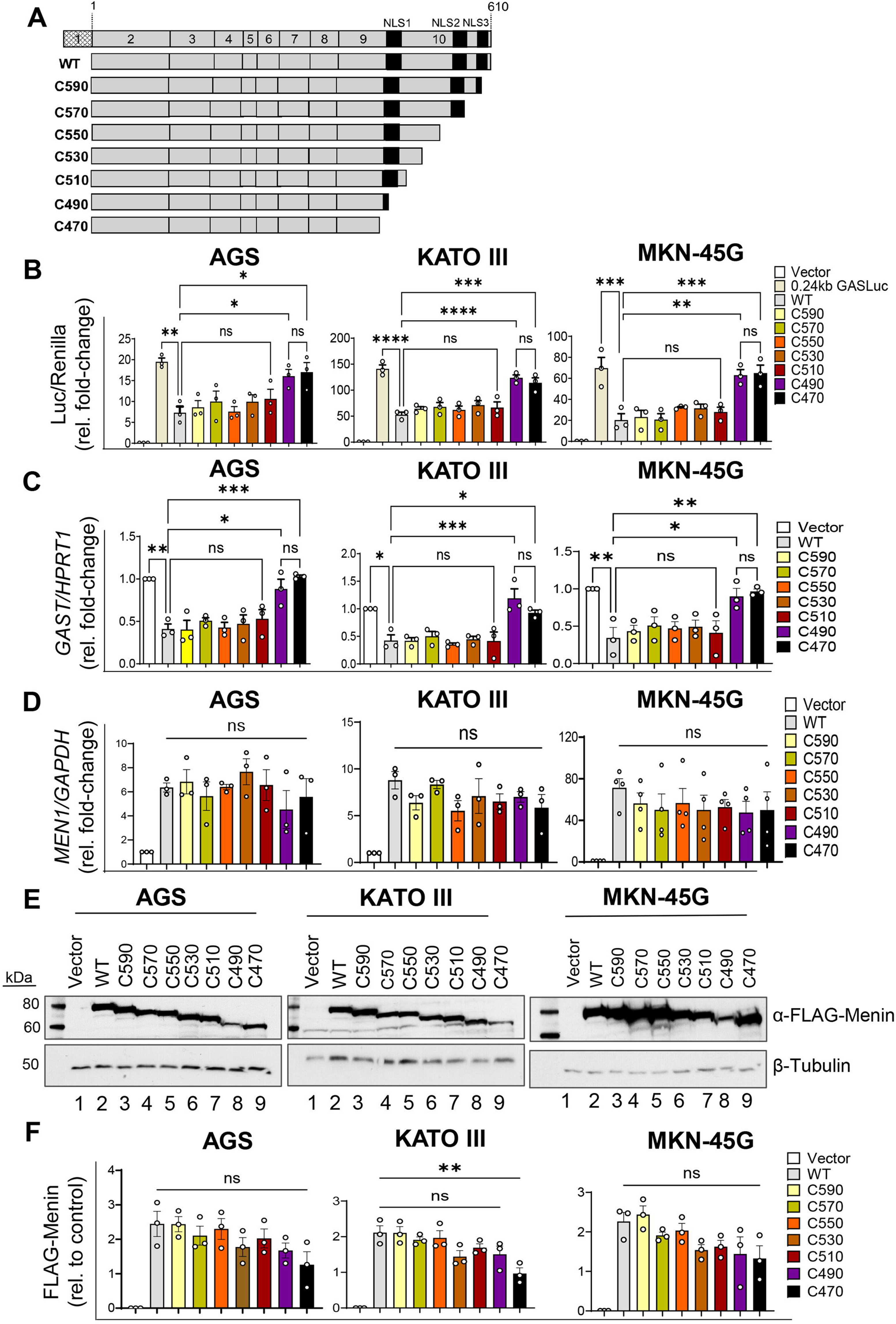
Ligand enrichment in DNETs correlates with loss of nuclear Menin localization. (A) Hematoxylin and eosin (H&E) staining of normal duodenum containing Brunner’s glands (nDUO-BG) and duodenal neuroendocrine tumor (DNET). Dashed boxes indicate regions shown at higher magnification. (B) Immunohistochemical staining for synaptophysin (SYP) confirming neuroendocrine differentiation in DNET. (C-D) Immunohistochemical staining for TGFα and EREG in tumor-associated Brunner’s glands (tBG) and DNET. Dashed boxes indicate tumor-gland interfaces. (E-F) Quantification of TGFα and EREG expression by H-score in nDUO-BG, tBG, and DNET. Data are mean ± SEM; ns, not significant; ****P < 0.0001. (G) EGFR immunostaining in nDUO-BG and DNET showing heterogeneous expression across tissues. (H) Menin immunostaining in nDUO-BG and DNET. (I) Representative FFPE DNET specimens showing cytoplasmic or near-absent Menin expression, accompanied by strong TGFα and EREG staining within tumor cells. (J) Quantification of Menin nuclear-to-cytoplasmic (N/C) ratio in nDUO-BG and DNET. Data are mean ± SEM; ****P < 0.0001. Images were taken at 100X, 200X and 400X. Scale bars: 100 μm (low magnification) and 50 μm (high magnification).

As expected [4,38], Menin protein expression was strong and nuclear in normal epithelium and BGs (**Fig. 2H, J**). In contrast, tumors exhibited reduced nuclear Menin, with 23% showing complete loss and the majority displaying cytoplasmic localization (**Fig. 2H, J)**. Although *MEN1* mutational status was unavailable for most cases due to poor DNA quality from archival FFPE specimens for sequencing, prior analysis demonstrates that loss of nuclear Menin can occur independently of *MEN1* mutational status [38], indicating alternative regulatory mechanisms. Consistent with this, tumors with reduced nuclear Menin displayed strong TGFα⁺ and EREG⁺ immunoreactivity (**Fig. 2I**), indicating an association between extracellular ligand enrichment and cytoplasmic Menin localization in DNET, correlating with signaling-dependent regulation rather than solely genetic loss of *MEN1*.

### EGFR signaling promotes cytoplasmic localization of Menin

To test whether EGFR signaling regulates Menin localization, we treated the gastrin-expressing KATO III, AGS, and MKN-45G cell lines with EREG. EREG stimulation showed no significant affect on total Menin protein levels. Next, we assessed whether EGFR activation regulates Menin subcellular localization and showed that wild-type Menin and constructs retaining at least NLS2 and NLS3 were distributed between nuclear and cytoplasmic compartments under basal conditions, whereas C490 and C470 mutants with a altered or complete loss of NLS1 were predominantly cytoplasmic. Similar results were demonstrated by immunofluorescence in AGS cells, indicating that intact NLS motifs are required for nuclear localization, while their loss promotes cytoplasmic retention.

Based on the differences between wild-type and NLS1–3 deleted mutants, subsequent analyses focused on wild-type, C490, and C470 constructs. To examine nuclear location in the absence of endogenous Menin, these constructs were expressed in the *Men1-null* mouse embryonic fibroblasts cells (MEF^Δ*Men1*^), and subcellular localization was assessed by IF microscopy. Upon EREG stimulation, nuclear FLAG–WT Menin levels decreased by approximately 75%, while cytoplasmic levels increased by ∼175% (**Fig. 3A, B**) [30]. In contrast, the NLS1-3 deleted C490 and C470 mutants remained predominantly cytoplasmic, with minimal or undetectable nuclear localization, and did not exhibit significant redistribution following EREG treatment (**Fig. 3A, B**). Proteasome inhibition with MG132 did not restore the nuclear localization of wild-type Menin and NLS1-3-deleted constructs (**Fig. 3A, B**), suggesting that loss of the nuclear import signals contributes to its cytoplasmic accumulation. To confirm these findings in a disease-relevant context, we repeated the IF analysis in AGS gastric carcinoma cells and observed similar EREG-induced reduction in nuclear wild-type Menin (**Fig. 3C, D**), consistent with impaired nuclear re-accumulation of Menin following EGFR activation in gastric cancer cells.

**Figure 3.**
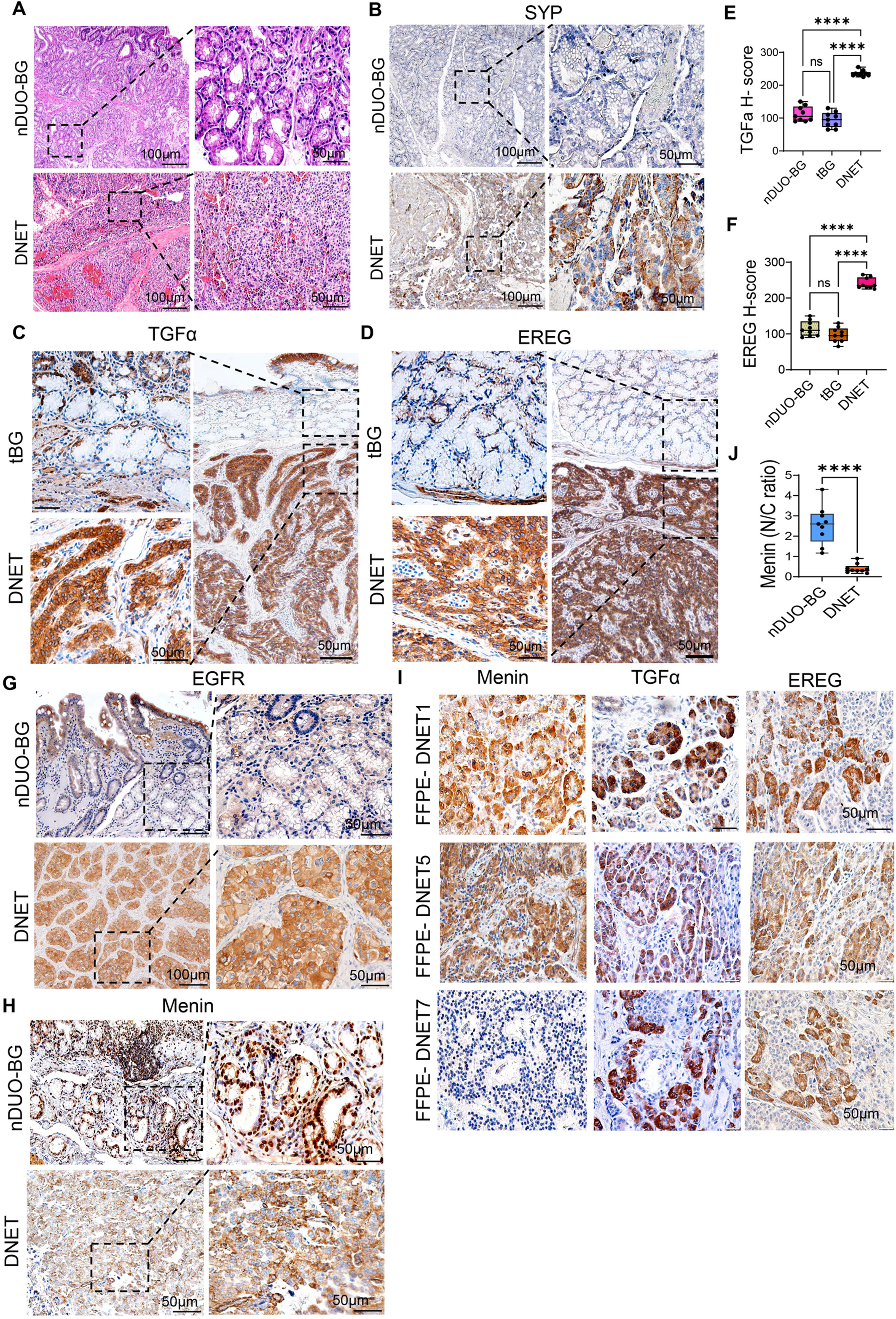
EGFR activation regulates Menin subcellular localization. (A, B) Representative IF images of FLAG-Menin–expressing MEF*^ΔMen1^* and AGS cells showing FLAG-WTMenin localization (white) under the indicated treatment conditions. Arrows indicate representative cells exhibiting cytoplasmic redistribution of Menin. Scale bar, 20 μm. (D, E) Quantification of FLAG-Menin subcellular localization in MEF*^ΔMen1^* and AGS cells. FLAG⁺ cells were scored per high-power field (HPF) and classified as nuclear (N) or cytoplasmic (C). Data are expressed as the percentage of nuclear (white) and cytoplasmic (gray/black) FLAG⁺ cells per HPF from ≥5 randomly selected fields per condition across ≥3 independent experiments. Images were analyzed at 400× Mag across the experiments. Data are presented as mean ± SEM. Statistical analysis was performed using two-way ANOVA with Sidak’s multiple-comparison test. ***P < 0.001. (E, F) Western blot analysis of nuclear and cytoplasmic protein extracts from AGS cells transfected with wild-typw Menin, C490, or C470 constructs with or without EREG and MG132. Lamin B1 (LMNB1) and β-tubulin served as nuclear and cytoplasmic loading controls, respectively. (G, H) Western blot analysis of nuclear and cytoplasmic protein extracts from KATO III cells transfected with wild-type Menin, C490, or C470 constructs following EREG and MG132 treatments. LMNB1 and β-tubulin served as nuclear and cytoplasmic loading controls, respectively. (I) Quantification of FLAG-Menin protein expression in AGS cells normalized to loading controls. (J) Quantification of FLAG-Menin protein expression in KATO III cells normalized to loading controls. Data are presented as mean ± SEM; Two-way ANOVA with Sidak’s post hoc test. ***P < 0.01; (n = 3 biological replicates).

To quantitatively assess Menin subcellular distribution, we performed biochemical fractionation followed by immunoblotting in AGS and KATO III cells. Under basal conditions, wild-type Menin was distributed approximately equally between nuclear and cytoplasmic fractions **(Fig. 3E, G, I, and J).** Upon EREG stimulation, nuclear wild-type Menin decreased by ∼70-75%, with a corresponding increase in cytoplasmic Menin despite minimal changes in total protein levels **(Fig. 3E, G, I, and J)**. Proteasome inhibition did not restore nuclear Menin but did increase cytoplasmic levels, indicating that EREG promotes nuclear export and turnover by limiting proteasome degradation. In contrast, since the C490 and C470 truncation mutants were largely restricted to the cytoplasm under basal conditions, with <1% detected in nuclear fractions, they were unresponsive to EREG stimulation **(Fig. 3F, H, I, and J)**. Proteasome inhibition increased cytoplasmic Menin expression in both AGS cells and KATO III cells. Together, these results indicate that EGFR signaling stimulates Menin’s translocation, prevents nuclear re-entry and promotes cytoplasmic retention.

### EGFR signaling derepress Menin-mediated *GAST* repression via nuclear localization

Prior studies have shown that EGFR signaling activates *GAST* expression, suggesting relief of Menin-mediated transcriptional repression [3,5,21]. Given the enrichment of EGFR ligands and reduced nuclear Menin observed in DNETs, *MEN1*-deficient models, and gastric NET cell lines, we investigated whether the Menin CTD, particularly its NLS motifs, is required for EREG-mediated regulation of *GAST*. Human gastric carcinoma cell lines (KATO III, AGS, and MKN-45G) were co-transfected with the 240 GASLuc reporter and wild-type Menin or C-terminal truncation mutants, followed by EREG stimulation for 6–8 h. Consistent with previous studies [5,21], EREG significantly increased *GAST* promoter activity. In cells expressing wild-type Menin, EREG partially relieved basal repression, resulting in a 1.8–2.2-fold increase in luciferase activity relative to untreated controls **(Fig. 4A–C)**. Similarly, truncation mutants retaining at least one intact NLS (C590–C510) showed a 1.6–2.0-fold induction, restoring promoter activity to near basal levels. In contrast, constructs lacking all three NLSs (C490–C470) failed to repress basal *GAST* expression and were unresponsive to EREG stimulation **(Fig. 4A–C).** Comparable results were observed at the transcript level: EREG increased endogenous *GAST* mRNA in cells expressing wild-type or NLS-retaining constructs, but not in cells expressing NLS-deficient mutants **(Fig. 4D–F).**

**Figure 4.**
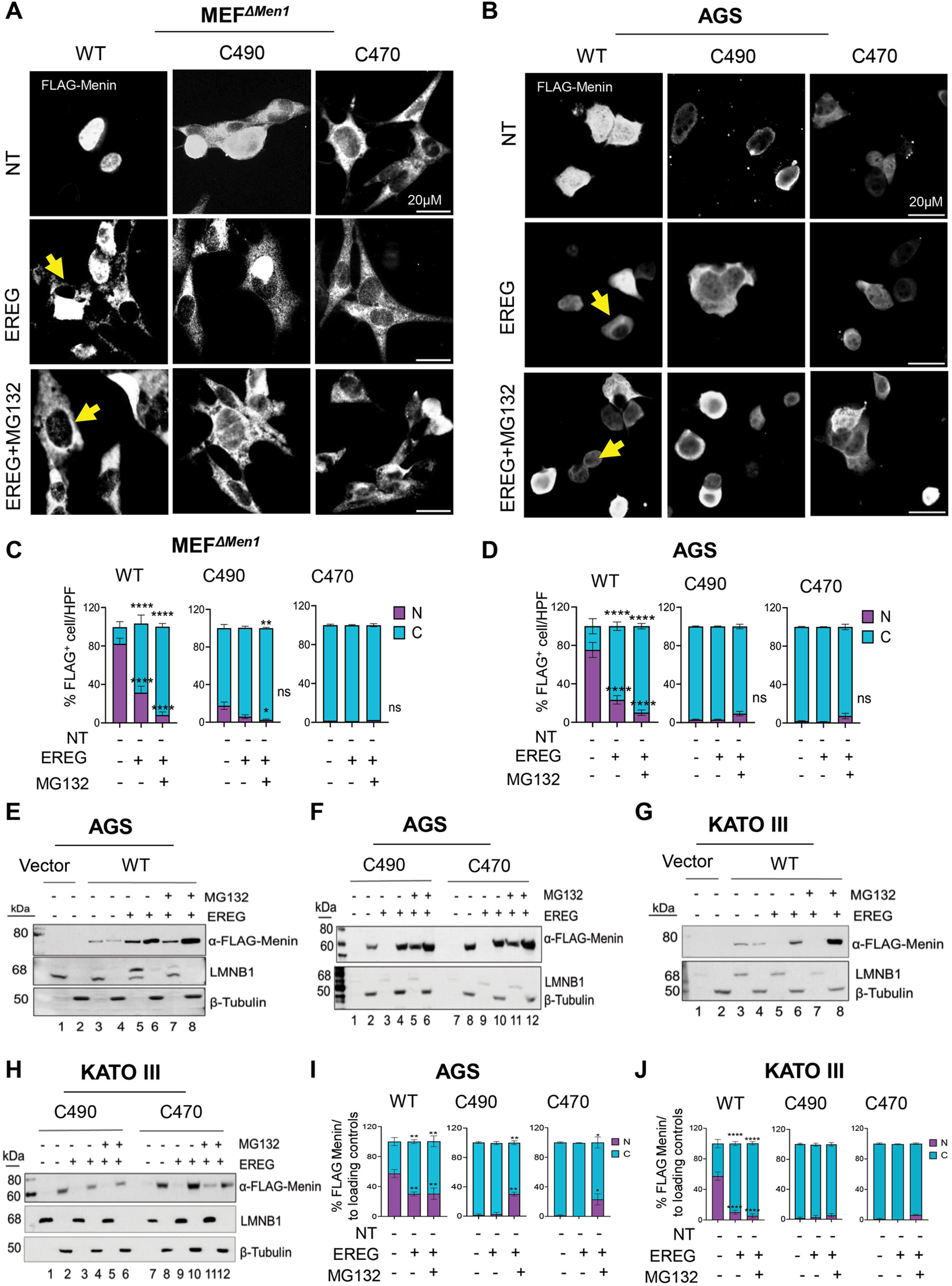
Menin CTD is required for EREG-mediated gastrin regulation. (A-C) Luciferase reporter assays in AGS, KATO III and MKN-45G cells expressing wild-type Menin or serial C-terminal truncation mutants (C590-C470). Cells were treated ± EREG, and luciferase activity was normalized to Renilla (Luc/Renilla, relative fold-change). (D-F) Quantitative RT-PCR analysis of endogenous *GAST* mRNA expression in KATO III, AGS, and MKN-45G cells expressing wild-type or truncated Menin constructs following ± EREG stimulation. Gene expression was normalized to *HPRT1* and expressed as relative fold-change. (G, H) *MEN1* mRNA levels in KATO III and MKN-45G cells following ± EREG treatment, normalized to *GAPDH*. Data are presented as mean ± SEM from n = 3-4 independent experiments performed in triplicate. Two-way ANOVA with Sidak’s post hoc test; *P < 0.05, **P < 0.01, ***P < 0.001, ****P < 0.0001; ns = not significant.

To determine whether these effects reflected altered *MEN1* transcription, we quantified *MEN1* mRNA levels following EREG stimulation in KATO III and MKN-45G cells. Quantitative RT-PCR revealed comparable *MEN1* transcript levels across all constructs in both cell lines **(Fig. 4G, H),** indicating that EGFR-mediated relief of repression does not arise from changes in *MEN1* expression. Together, these data define a spatially organized and mechanistically coherent framework for EGFR–Menin control of *GAST* expression in the duodenum. The integrity of the Menin CTD is required to maintain basal repression of GAST; however, EREG-induced activation occurs despite intact NLS1–3 and without changes in *MEN1* transcript abundance. These findings exclude transcriptional downregulation as a primary mechanism and instead support a model in which EGFR signaling regulates *GAST* expression through post-translational control of nuclear Menin.

### Extracellular signaling induces Menin phosphorylation at Ser487 within the NLS1 region

The CTD of Menin contains five predicted phosphorylation sites [39], including Ser487 in the highly conserved NLS1 region (**Fig. 5A, B**), suggesting possible post-translational regulation. The sequence surrounding Ser487 contains a basic residue-rich motif, including an Arg-Arg-Glu-Ser (RRES) sequence that matches consensus phosphorylation motifs recognized by the AGC family of kinases (i.e., protein kinase A, G, and C). Forskolin (FSK) is a direct activator of adenylyl cyclase, cAMP and PKA pathway. In some instances, 12-O-tetradecanoylphorbol-13-acetate (TPA), a potent inducer of PKC can indirectly activate PKA[40] (**Fig 5A, B**). To identify candidate upstream regulators, we reviewed an in-silico kinase prediction website that ranked Ser487 as a high-confidence phosphorylation site targeted by multiple AGC family kinases, including Protein Kinase, X-Linked (PRKX) and PKA. Both PKA and PRKX are serine/threonine kinases implicated in cAMP-dependent signalling pathways that regulate proliferation and differentiation [41,42]. These predictions were used to guide hypothesis generation and contextualize downstream experimental analyses.

**Figure 5.**
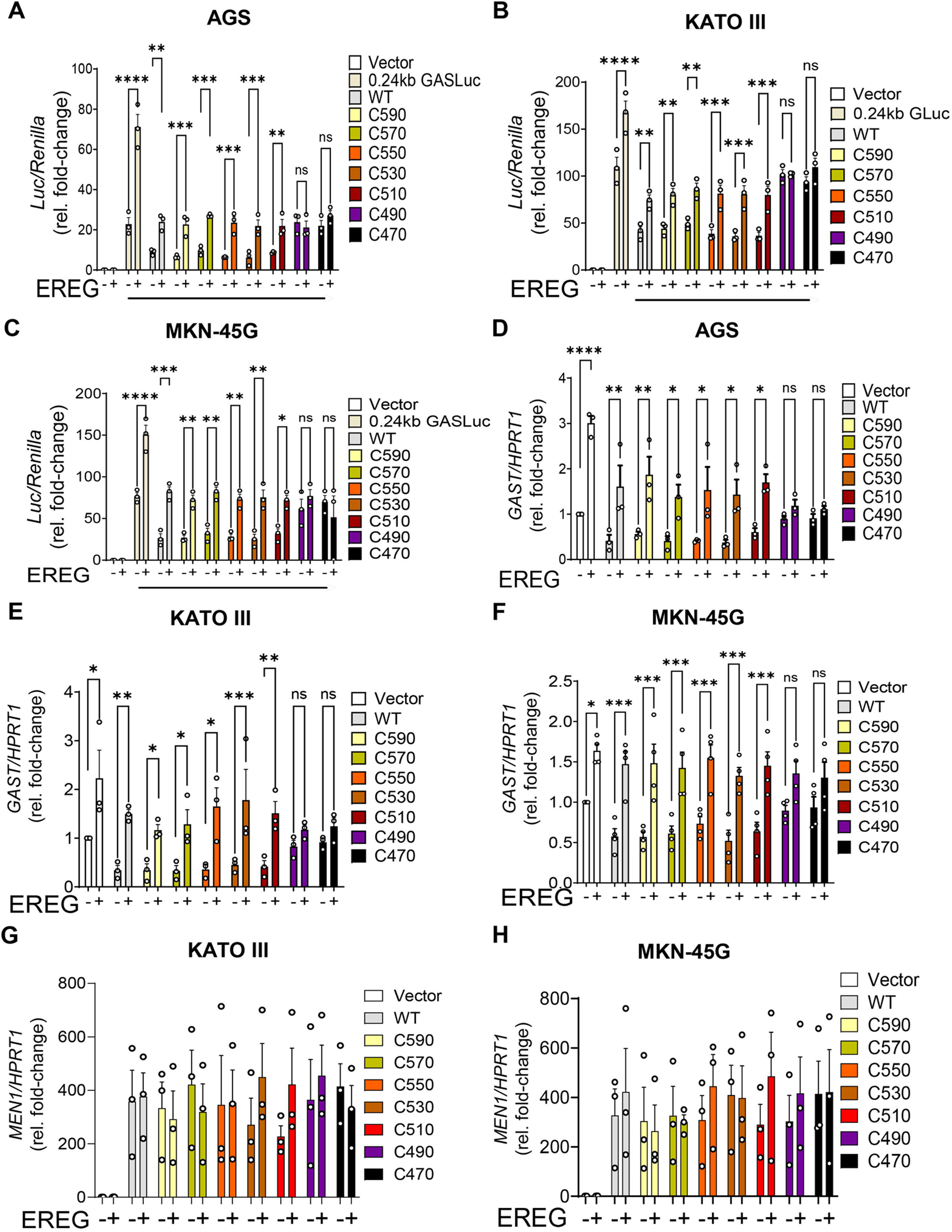
Extracellular signaling induces phosphorylation of Menin at Ser487. (A) Multiple sequence alignment of the Menin C-terminal region from the indicated vertebrate species showing strong conservation of a basic residue–rich motif encompassing Ser487. Conserved basic residues and Ser487 are highlighted. (B) Schematic of human Menin illustrating the position of Ser487 within NLS1. The expanded sequence highlights Ser487 and surrounding basic residues; constructs used in this study. (C) Immunoblot analysis of AGS cells expressing FLAG-tagged wild-type Menin or Ser487 mutants (S487A, S487D) following treatment with EREG, FSK, or TPA. Whole-cell lysates were probed with antibodies against phospho-Ser487 Menin, FLAG-Menin, and GAPDH. (D, E) Immunoblot analysis of MKN-45G and KATO III cells expressing wild-type Menin following stimulation with EREG, FSK, or TPA. Blots were probed for phospho-Ser487 Menin, FLAG-Menin, and β-tubulin. (F, H) Quantification of phospho-Ser487 Menin in AGS, KATO III and MKN-45G cells. (I) Immunoblot analysis of AGS cells examining activation of cAMP and EGFR downstream kinases under the indicated conditions. (J) Densitometric quantification of signalling outputs shown in (I), expressed as fold change relative to vector control. (K) Time-course of Ser487 phosphorylation in AGS cells stimulated with TPA in the presence of kinase inhibitors; MEK inhibitor (U0126), AKT inhibitor (MK-2206), PKC inhibitor (Gö6983), or combined MEK+AKT inhibition. (L) Quantification of Ser487 phosphorylation kinetics following TPA stimulation with the indicated inhibitors. (M) Area-under-the-curve (AUC) analysis of phosphorylation in (L). Data are presented as mean ± SEM; individual data points represent independent biological replicates (n = 3). Statistical significance was determined by one-way ANOVA with Tukey’s multiple-comparison test (*P < 0.05; **P < 0.01; ****P < 0.0001; ns, not significant).

To directly assess Ser487 phosphorylation, we used a previously generated phospho-specific monoclonal antibody recognizing phospho-Ser487 Menin [24]. Antibody specificity was validated by immunoblotting whole-cell lysates expressing FLAG-tagged wild-type Menin or Ser487 mutant constructs (S487A and S487D). A phospho-dependent signal was detected in wild-type Menin but was absent in both mutants under identical conditions (**Fig. 5C**). Signal specificity was further supported by increased immunoreactivity following pathway stimulation with EREG, FSK, or TPA (**Fig. 5C**). Total Menin expression was confirmed using anti-FLAG antibody, and β-tubulin served as a loading control. Using this validated antibody, we next examined Ser487 phosphorylation in AGS, MKN-45G, and KATO III gastric cancer cell lines. Under basal conditions, wild-type Menin exhibited detectable Ser487 phosphorylation across all three cell lines **(Fig. 5C–H).** Stimulation with EREG, FSK, or TPA significantly increased Ser487 phosphorylation **(Fig. 5C–H).** These findings establish Ser487 as a dynamically regulated phosphorylation site responsive to extracellular signaling.

Given that EREG and TPA activate multiple downstream signaling pathways, including MAPK and AKT, we assessed pathway-specific kinase activation. Notably, NLS1 contains a weak ERK consensus motif (Pro-X-S/T-Pro) [43], suggesting that MAPK signaling might contribute to phosphorylation at Ser487. Additional inspection of the sequence context surrounding Ser487 (RRRGPRRESKP) revealed a motif compatible with, but not strictly canonical for an AKT phosphorylation site (RXRXXS/T) [44]. To delineate pathway-specific contributions, we examined downstream kinase activation in AGS cells. EREG robustly induced phosphorylation of AKT and ERK1/2, consistent with activation of canonical EGFR signaling. Total ERK levels showed a modest non-significant decrease (**Fig. 5I, J**). In contrast, FSK selectively increased CREB phosphorylation without appreciable changes in PKA abundance or phosphorylation of PKA substrates (**Fig. 5I, J**), indicating activation of cAMP-dependent signaling. Notably, TPA elicited strong activation of ERK1/2 and CREB, with concurrent AKT phosphorylation. Although total ERK levels trended downward relative to tubulin, this change was not statistically significant (**Fig. 5I, J**), suggesting potential crosstalk among PKC-dependent signaling pathways (**Fig. 5I, J**).

Next, we determined whether Ser487 phosphorylation functions as a dynamically regulated node downstream of convergent extracellular signaling pathways. Time-course analysis of pS487 in AGS cells following TPA stimulation revealed minimal basal phosphorylation, followed by a rapid increase detectable by 5 min that peaked at 30 min (**Fig. 5K, L**). Phosphorylation declined by 60 minutes and subsequently increased again at 120 min but at a lower amplitude, indicating transient attenuation of the signal. To define pathway dependency, AGS cells were pretreated with selective inhibitors targeting AKT (MK-2206), MEK1/2 (U0126), or PKC (Gö6983) prior to TPA stimulation. Inhibition of AKT or MEK partially attenuated Ser487 phosphorylation, whereas PKC inhibition significantly reduced phosphorylation **(Fig. 5K–M)**, indicating PKC as a dominant upstream regulator under PKC-driven conditions. Combined inhibition of AKT and MEK further reduced phosphorylation to levels comparable to PKC blockade **(Fig. 5K–M)**, supporting a model in which PKC functions upstream of multiple kinase branches that converge on Ser487. Loading controls remained unchanged, indicating that the observed effects reflect altered phosphorylation rather than changes in Menin abundance.

### Phosphorylation of Ser487 Abrogates Menin-Mediated Repression of *GAST*

Previous work in insulinoma cells showed that phosphorylation of Menin at Ser487 impairs its ability to repress insulin transcription [24]. To determine whether this regulatory mechanism extends to gastrin-expressing cells, we assessed the function of Ser487 mutation on endogenous *GAST* gene expression. AGS, MKN-45G, and KATO III cells were transfected with wild-type, phospho-null (S487A), or phospho-mimetic (S487D) Menin constructs. Under basal conditions, S487A Menin did not exhibit significant suppression of endogenous *GAST* mRNA expression relative to wild type, whereas the S487D mutant induced rather than repressed gastrin mRNA **(Fig. 6A–C)**, indicating that phosphorylation at Ser487 disrupts Menin-mediated transcriptional repression. To determine whether these effects did not reflect altered Menin express, *MEN1* mRNA levels were measured following overexpression of wild-type and mutant constructs. and showed no difference in construct expression **(Fig. 6D, E).** Importantly, *MEN1* mRNA levels did not differ significantly between wild-type, S487A, and S487D constructs, despite lower protein levels observed for S487A (**Fig. 5C**).

**Figure 6.**
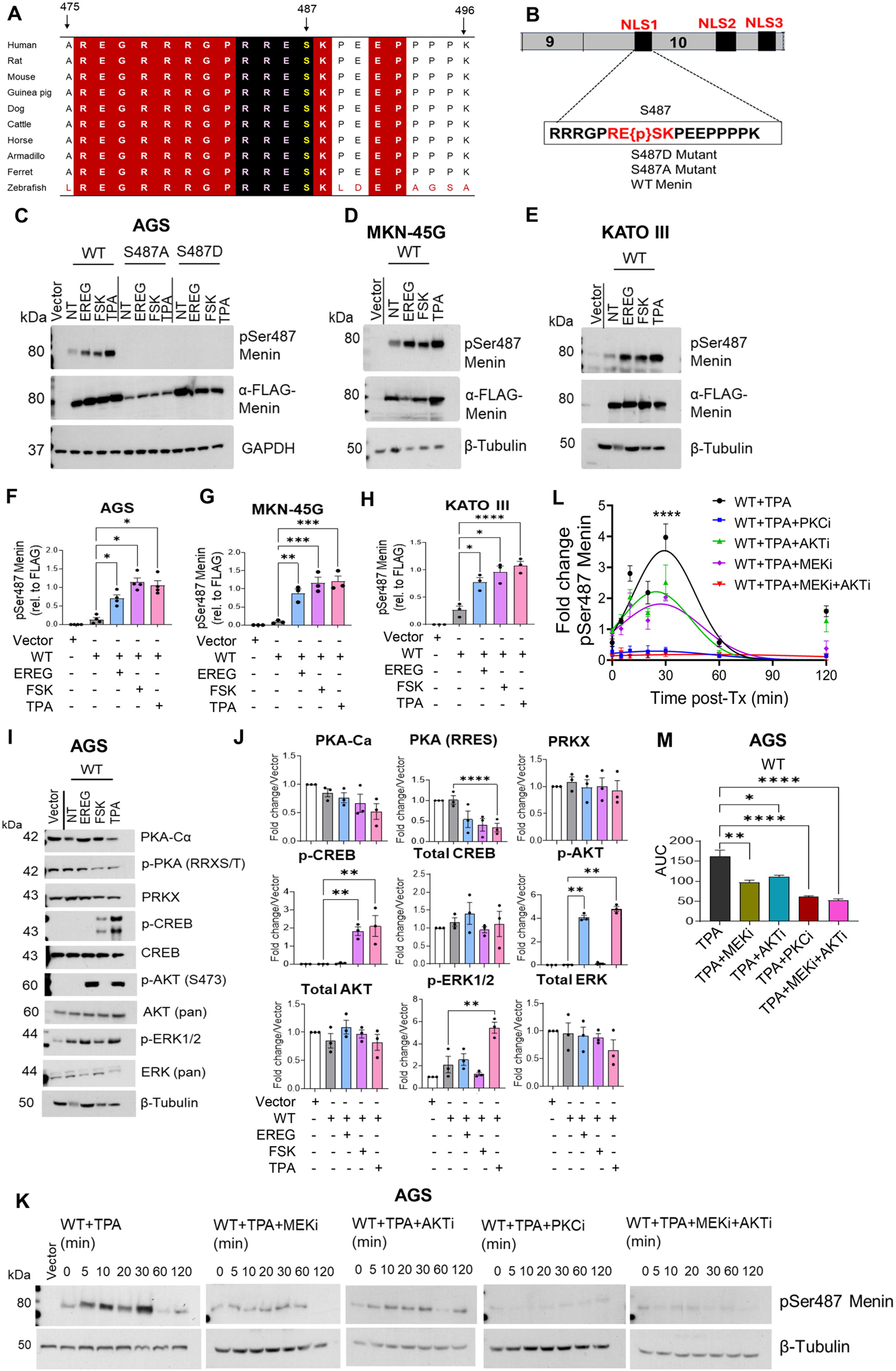
Phosphorylation of Menin at Ser487 impairs repression of *GAST* expression. (A–C) Quantitative RT–PCR analysis of endogenous GAST mRNA levels in MKN-45G, AGS, and KATO III cells transfected with wild-type, S487A, or S487D Menin constructs. (D, E) Quantification of *MEN1* mRNA expression following overexpression of wild-type and mutant Menin constructs in MKN-45G and AGS cells. Data represent mean ± SEM from three independent experiments. One-way ANOVA with Tukey’s multiple-comparison test (**P < 0.01; ***P < 0.001; ****P < 0.0001). (F–I) Effect of EREG or FSK treatment on endogenous *GAST* mRNA expression in KATO III, MKN-45G, and AGS cells expressing wild-type or mutant Menin constructs. qRT–PCR data were normalized to *HPRT1*, as indicated, and expressed relative to vector control. (J, K) *GAST* promoter activity measured using the 240 *GASLuc* reporter in AGS and KATO III cells co-transfected with wild-type or mutant Menin constructs and treated with TPA. (L) AGS cells transfected with the 240 GASLuc reporter were treated with FSK in the presence or absence of kinase inhibitors following 1 h pretreatment: H89 (10 μM), MK-2206 (5 μM), U0126 (10 μM), and Gö6983 (5 μM). (M) AGS cells transfected with *GASLuc* were treated with TPA in the presence of the indicated inhibitors following 1 h pretreatment. Data represent mean ± SEM (n = 3 independent experiments). Statistical analysis was performed using one-way or two-way ANOVA with Tukey’s or Sidak’s multiple-comparison test, as indicated (*P < 0.05; **P < 0.01; ****P < 0.0001).

To assess the effect of extracellular signaling on Menin-dependent repression of endogenous *GAST* after phosphorylation of Menin at S487, KATO III, AGS, and MKN-45G cells transfected with wild-type, S487A, or S487D Menin were treated with EREG or FSK. EREG significantly induced *GAST* mRNA levels. However, the phospho-deficient S487A mutant remained resistant to signal-induced induction of *GAST* **(Fig. 6F, G).** In contrast, the phospho-mimetic S487D mutant showed no repressive activity at baseline and failed to respond to FSK stimulation (**Fig. 6F, G**), consistent with constitutive loss of Menin function resulting in constitutive activation of endogenous *GAST*. Similar patterns were observed following FSK treatment of KATO III and MKN-45G cells, where wild-type Menin repression was relieved by ligand stimulation, while the S487A mutant remained resistant to derepression and the S487D mutant showed constitutive *GAST* induction **(Fig. 6H, I).** To determine whether PKC signaling produced a similar effect, KATO III and AGS cells were co-transfected with the 240 *GASLuc* reporter together with wild-type or mutant Menin constructs and stimulated with TPA. Consistent with the effects of FSK and EREG, TPA significantly attenuated wild-type Menin-mediated repression of the *GAST* promoter (**Fig. 6J, K**). The S487A mutant remained resistant to signal-induced derepression, whereas the S487D mutant failed to repress basal promoter activity, showed constitutive activation and remained unresponsive to further induction by TPA (**Fig. 6J, K**). Together, These results demonstrated thatSer487 phosphorylation is a critical determinant of Menin’s transcriptional repressor activity that links cAMP- and EGFR-dependent signaling pathways to Menin inactivation in *GAST*-expressing cells.

To further define the signaling pathways required for stimulus-induced derepression of *GAST*, we tested inhibitors of the various signaling pathways in AGS cells since they showed the strongest response to extracellular stimulation. AGS cells were co-transfected with the *240 GASLuc* reporter together with wild-type Menin. Because FSK and TPA produced the most robust induction of *GAST* expression, cells were pretreated for 1 h with the selective kinase inhibitors and then stimulated with FSK or TPA at the indicated concentrations. FSK significantly increased *GAST* promoter activity, and this effect was attenuated by inhibition of PKA, AKT, MEK, or PKC, with the strongest suppression observed when MEK and AKT were inhibited simultaneously **(Fig. 6L).** Similarly, TPA-induced activation of the *GAST* promoter was significantly reduced below baseline by PKC inhibition and partially attenuated by inhibition of AKT or MEK, whereas PKA inhibition had minimal effect **(Fig. 6M).** Combined inhibition of MEK and AKT further suppressed promoter activity, consistent with cooperative signaling downstream of PKC. Collectively, these results demonstrated that extracellular signaling relieves Menin-dependent repression of *GAST* through convergent kinase pathways.

### Ser487 Phosphorylation Disrupts Nuclear Localization and Stability

We previously showed in enteric glia that gastrin-Cckbr signalling induces Menin nuclear export via PKA [30]. To determine whether phosphorylation at Ser487 mediates a similar process in Menin-null and gastric adenocarcinoma cells, wild-type, phospho-null (S487A), and phospho-mimetic (S487D) Menin constructs were expressed in MEF^Δ*Men1*^ and KATO III cells and analysed by IF. In both cell lines, wild-type and S487A Menin localized predominantly to the nucleus and perinuclear compartment **(Fig. 7A-C)**, consistent with earlier reports [25,30]. However, the S487D mutant was significantly localized to the cytoplasm (**Fig. 7A-C**). To validate these observations biochemically, subcellular fractionation followed by immunoblotting was performed in KATO III and AGS gastric carcinoma cells. Under basal conditions, wild-type Menin was predominantly localized to the nucleus, with ∼70–80% detected in the nuclear fraction **(Fig. 7D–G).** The phospho-deficient S487A mutant displayed a more equal nuclear–cytoplasmic distribution, whereas the phospho-mimetic S487D mutant was largely restricted to the cytoplasmic fraction, with ∼80% detected outside the nucleus in all cells **(Fig. 7D–G).** These results indicate that phosphorylation at Ser487 alters Menin subcellular localization.

**Figure 7.**
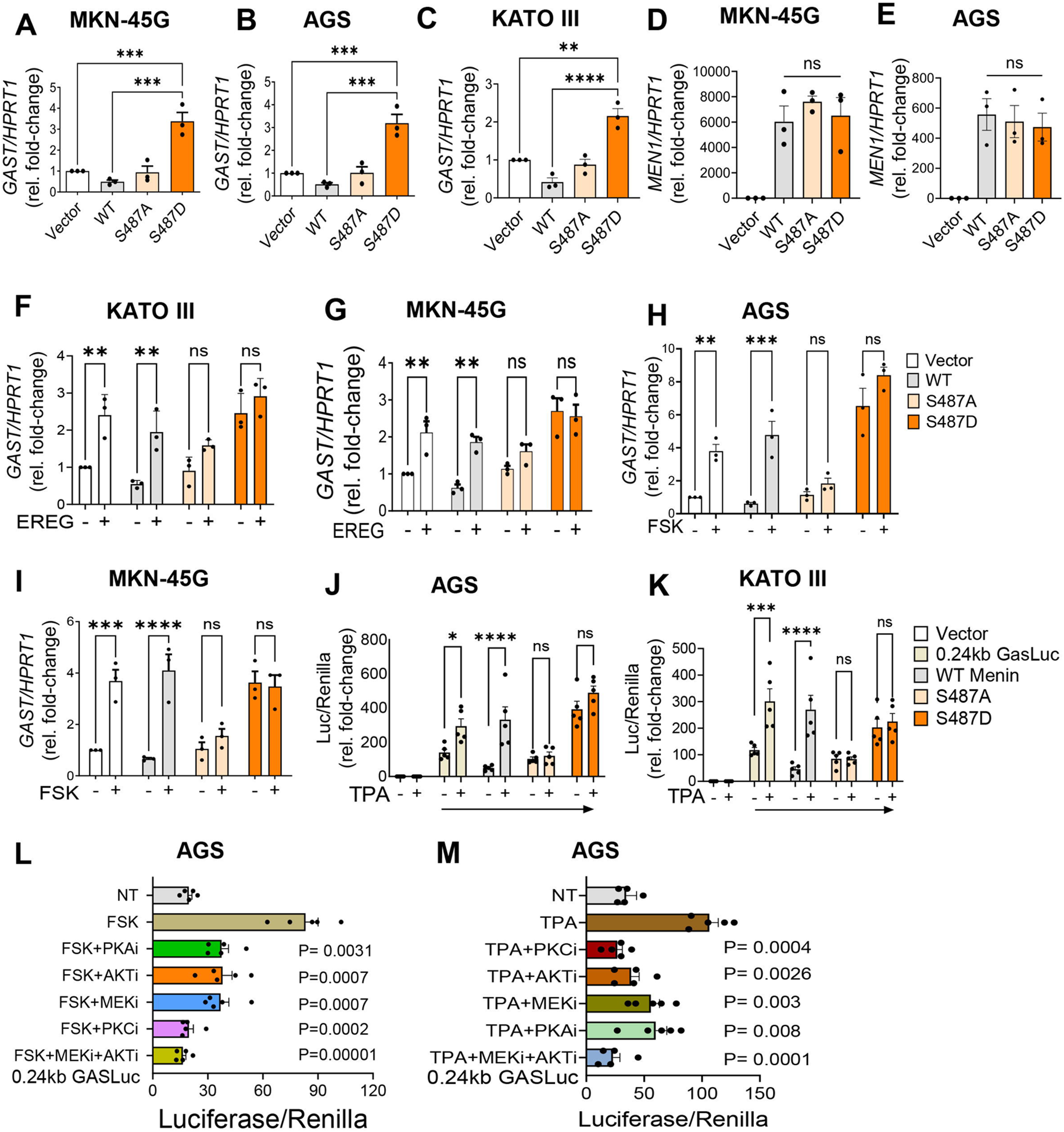
Ser487 phosphorylation regulates Menin subcellular localization and signal-dependent nuclear export. (A–C) Representative IF images and quantification of FLAG-Menin localization in MEF*^ΔMen1^* and KATO III cells expressing wild-type Menin, phospho-null (S487A), or phospho-mimetic (S487D) constructs. Cells were treated with forskolin (FSK) to activate cAMP/PKA signaling or TPA to activate PKC signaling. FLAG-Menin is shown in white. Arrows indicate cells exhibiting cytoplasmic Menin accumulation. Scale bar, 20 μm. (B, C) Quantification of nuclear and cytoplasmic FLAG-Menin localization expressed as the percentage of FLAG-positive cells per high-power field (HPF). Data represent mean ± SEM from at least five randomly selected fields and analyzed per condition at 400× magnification. Statistical significance was determined relative to untreated controls. (D, E) Subcellular fractionation and immunoblot analysis of FLAG-Menin localization in KATO III and AGS cells expressing wild-type, S487A, or S487D constructs following pathway activation (FSK or TPA as indicated). Nuclear and cytoplasmic fractions were analyzed by immunoblot using anti-FLAG antibodies. Histone H3 and β-tubulin served as nuclear and cytoplasmic loading controls, respectively. Representative blots are shown with both high- and low-exposure FLAG-Menin signals. (F, G) Densitometric quantification of FLAG-Menin distribution between nuclear and cytoplasmic fractions in KATO III and AGS cells, normalized to the corresponding loading controls. (H, I) Quantification of nuclear and cytoplasmic Menin localization expressed as the percentage of FLAG-positive cells per HPF. Data represent mean ± SEM from at least five randomly selected fields analyzed per condition at 400× magnification. Scale bars: 20 µm. (J) IF analysis of Menin localization in AGS and MEF*^ΔMen1^* cells under basal conditions and following treatment with TPA, the nuclear export inhibitor leptomycin B (LMB), or the proteasome inhibitor MG132, as indicated. Menin is shown in white. Data are presented as mean ± SEM from ≥3 independent experiments. Statistical analysis was performed using one-way or two-way ANOVA with Tukey’s multiple-comparison test, as indicated (*P < 0.05, **P < 0.01, ***P < 0.001, ****P < 0.0001; ns, not significant).

We next examined the effect of extracellular signaling on Menin localization following FSK or TPA treatment. FSK significantly altered the subcellular distribution of wild-type Menin in KATO III cells, reducing the nuclear fraction to ∼40% (**Fig. 7D, F**). The phospho-deficient S487A mutant distributed equally between the nuclear and cytoplasm but was largely resistant to FSK-induced redistribution, whereas the phospho-mimetic S487D mutant was constitutively enriched in the cytoplasmic fraction and showed no further change following stimulation (**Fig. 7D, F**). These results are consistent with phosphorylation at Ser487 impairing nuclear retention of Menin. Similar effects were observed following TPA treatment in AGS cells (**Fig. 7E, G).**To test whether this redistribution reflected chromosome region maintenance 1 (CRM1) -dependent nuclear export, MEF*^ΔMen1^* and gastric carcinoma cells (AGS) expressing wild-type Menin were treated with the CRM1 nuclear export inhibitor leptomycin B (LMB) and analyzed by IF. We used TPA to stimulate nuclear export since it was the most potent inducer of Menin translocation. LMB decreased nuclear export of Menin reducing its accumulation in the cytoplasm (**Fig. 7H–J**). These findings indicate that inducible phosphorylation promotes nuclear export of Menin, rather than simply preventing nuclear import. MG132 did not restore nuclear localization but instead allowed cytoplasmic Menin pools to accumulate, suggesting that the WT phosphorylated menin is subject to proteasomal degradation (**Fig. 7H-J**). Together, these data indicate that Ser487 phosphorylation promotes signal-dependent redistribution of Menin to the cytoplasm as a mechanism to derepress target genes like *GAST*.

## DISCUSSION

The molecular mechanisms by which extracellular growth factor signaling interfaces with lineage-specific tumor suppressors to modulate neuroendocrine tumor biology remain poorly defined. EGFR signaling remains a common regulator of tumor proliferation and survival, and its dysregulation has been widely implicated across multiple solid tumors [45–47]. Although EGFR-targeted therapies have demonstrated clinical benefit in malignancies such as colorectal cancer [48,49], their application in GEP-NETs has been limited, despite early evidence that these foregut tumors express both the receptor and ligand [37,50,51]. Pharmacologic inhibition of the EGFR and related growth factor pathways including insulin-like growth factor (IGF) and PI3K/AKT/mTOR signaling has largely resulted in disease stabilization rather than objective tumor regression [52–54], reflecting intertumoral heterogeneity, limited biomarker stratification, and compensatory signaling through redundant pathways. Consistent with this, mTOR inhibition with everolimus primarily prolongs progression-free survival without inducing substantial tumor shrinkage [54], underscoring the predominantly cytostatic nature of targeting these pathways. Thus, the lack of clinical response may be related to the need to better subtype the tumors [50]. These observations suggest that extracellular signaling in GEP-NETs does not function through linear, receptor-centric mechanisms alone, but instead converges on downstream regulatory hubs that integrate environmental cues to control tumor behavior.

Our findings establish a mechanistic link between ligand-driven EGFR signaling and functional inactivation of Menin that occurs independently of *MEN1* genetic loss. Rather than altering *MEN1* gene expression, we show that extracellular signaling regulated Menin post-translationally by phosphorylation of its CTD subsequently inducing its subcellular re-localization and cytoplasmic stability. These findings have important implications for MEN1-associated tumorigenesis, as clinicopathologic studies show that early duodenal lesions frequently retain Menin protein expression and a wild-type *MEN1* allele [11,55]. Our studies support an explanation by demonstrating that extracellular signaling pathways remove Menin from the nucleus which functionally inactivates Menin repression. NETs that inactivate Menin’s CTD by frameshift or missense mutations that disrupt the NLSs may also induce tumorigenesis without detectable loss of Menin protein. Persistent Menin protein in the cytoplasm, may provide evidence supporting this mechanism. The spatial enrichment of EGFR ligands, including TGFα and EREG, within DNETs further supports a ligand-driven signaling paradigm. Notably, this occurs despite constitutive EGFR expression. Indeed, we found that an increase in some tumor cells within the DNET expressed increased levels of both EREG and TGFa ligands, which may explain the heterogeneity of tumor responsiveness to EGFR inhibition. In the growth factor-rich microenvironment such as the proximal duodenum, even low receptor levels may be sufficient to sustain signaling capable of modulating Menin localization and function, providing a biological explanation for the continued relevance of EGFR pathways in GEP-NETs despite minimal therapeutic responses.

Our data indicate that multiple pathways activated by the EGFR, such as PKC and AKT and to a lesser extent MEK with some crosstalk to cAMP/PKA signaling, converge on a common regulatory node at Ser487, suggesting that inhibition of EGFR alone may be insufficient to restore Menin tumor-suppressor function. Instead, therapeutic strategies that preserve or restore nuclear Menin localization-such as targeting CRM1-dependent nuclear export or disrupting kinase convergence at Ser487-may provide a more effective approach.

Crosstalk of EGFR, MAPK, and cAMP-dependent pathways on promoter-regulatory proteins is a well-established mechanism for integrating extracellular cues with transcriptional control [21,22,32]. At the mechanistic level, our data identify phosphorylation of Ser487 within the NLS1 region as the central signaling node linking extracellular kinase activity to Menin regulation. Ser487 phosphorylation was induced by EREG, FSK, and TPA, demonstrating crosstalk of EGFR, cAMP/PKA, and PKC pathways at this site and positioning it as a nodal integration point rather than a component of a linear cascade. Functionally, this modification limits nuclear Menin availability through a CRM1-dependent export mechanism, as evidenced by partial restoration of nuclear localization following LMB treatment. Notably, the phospho-null S487A mutant remains nuclear, suggesting that phosphorylation at this residue cooperates with nuclear export signals to facilitate cytoplasmic redistribution, although the contribution of proximal export motifs has not yet been tested. In contrast, the phospho-mimetic S487D mutant fails to re-enter the nucleus and is not subject to degradation, raising the possibility that phosphorylation at this site may uncouple nuclear export from ubiquitin-mediated turnover. These findings define Ser487 phosphorylation as a transport-coupled regulatory switch that links extracellular signaling to Menin redistribution.

Consequences of this redistribution are most evident at the level of transcriptional control. Menin acts as a scaffold linking transcription factors such as JunD to chromatin-modifying complexes, including mSin3A-HDAC and MLL-associated regulators [3,56,57], and repression of the *GAST* expression is governed by promoter-proximal AP-1 elements [5]. Reduced nuclear Menin is expected to decrease promoter occupancy and impair recruitment of co-repressor complexes, thereby facilitating transcriptional activation in response to extracellular signaling. In this context, signaling-dependent control of Menin localization provides a direct mechanism for derepression of the *GAST* promoter without requiring changes in *MEN1* transcription.

The functional impact of Ser487 phosphorylation appears to be tissue dependent, which may explain differences between pancreatic β-cell function and gastric/duodenal NET tumorigenesis evaluated here. In normal β cells, this modification primarily affects transcriptional complex activity [24], whereas in gastroenteric NETs, it predominantly regulates nuclear localization. These differences likely reflect cell-type-specific kinase hierarchies, nuclear transport dynamics, and chromatin interactions, as well as distinct transcriptional partners engaged by Menin across lineages. In gastrointestinal cells, extracellular cues have been shown to promote both nuclear export and proteasome-dependent turnover of Menin [30], consistent with the transport-based mechanism defined here. Together, these findings provide a mechanistic explanation for the long-recognized “*MEN1* paradox,” in which early neuroendocrine lesions retain Menin expression yet exhibit reduced tumor-suppressor activity. We propose that extracellular signaling acts as an early functional hit by reducing the pool of nuclear, transcriptionally active Menin. In growth factor-rich niches such as BGs, sustained activation of EGFR-, PKC-, and cAMP-dependent pathways may lower the threshold for *GAST* derepression and clonal expansion, potentially preceding or cooperating with genetic loss.

In summary, this study identifies the C-terminal nuclear localization module of Menin as a signaling-sensitive domain that links extracellular kinase activity to transcriptional control of the *GAST* gene. Phosphorylation of Ser487 within NLS1 functions as switch that restricts nuclear Menin and disrupts promoter-associated repression. This mechanism provides a molecular framework for how growth factor-dependent signaling can override *MEN1* tumor-suppressor function without requiring immediate genetic loss and highlights a transport-dependent regulatory axis through which microenvironmental signals modulate Menin activity. These findings suggest that restoring nuclear Menin localization may represent a therapeutic strategy in MEN1-intact tumors especially in the duodenal lumen where they are exposed to sustained extracellular signaling.

## DECLARATIONS

### Patient consent for publication

Not applicable

## Acknowledgements

We thank Dr. Suleiman Sheriff and members of the Merchant laboratory for their assistance in generating the Menin C-terminal domain constructs. We acknowledge the University of Arizona Tissue Repository for providing biospecimens used in this study. We also thank Dr. Tobias Else, Dr. Michelle Kim, and the Biorepository and Pathology Core at the Icahn School of Medicine at Mount Sinai for their assistance with specimen collection and processing, and we gratefully acknowledge the biorepository participants for their contribution to this research. Graphical abstract Created in BioRender. Elvis--offiah, U. (2026) https://BioRender.com/uan4a7d

## Authors contributions

Uloma B. Elvis-Offiah: Data curation, formal analysis, supervision, validation, investigation, visualization, methodology, writing-original draft, writing-review and editing. Ziyu Wen: Resources, writing-review and editing. Xianxin Hua: Resources, writing-review and editing. Juanita L. Merchant: Conceptualization, resources, supervision, project administration, writing-review and editing.

## Supplementary data

Supplementary Data are available at NAR online.

## Conflict of interest

Authors declare no conflict of interest

## Funding

This was supported by PHS grant R01 DK45729 to JLM; P30CA062203 UACC Shared Resources and TARGHETS biospecimen core

## Data availability

All data generated or analyzed during this study are included in this article and its Supplementary Data. Source data underlying the figures are available online at NAR.

## Notes

### Competing Interest Statement

The authors have declared no competing interest.

## REFERENCES

1. Cingam SR, Botejue M, Hoilat GJ et al., StatPearls. StatPearls Publishing Copyright © 2025, StatPearls Publishing LLC., Treasure Island (FL),2025.

2. Jensen RT and Ito T. In Feingold, K. R., Anawalt, B., Boyce, A., Chrousos, G., de Herder, W. W., Dhatariya, K., Dungan, K., Hershman, J. M., Hofland, J., Kalra, S. et al. (eds.), Endotext. MDText.com, Inc. Copyright © 2000-2022, MDText.com, Inc., South Dartmouth (MA),2000.

3. Agarwal SK, Guru SC, Heppner C et al. Menin interacts with the AP1 transcription factor JunD and represses JunD-activated transcription. Cell. 1999; 96: 143–152.

4. Chandrasekharappa SC, Guru SC, Manickam P et al. Positional cloning of the gene for multiple endocrine neoplasia-type 1. Science. 1997; 276: 404–407.

5. Mensah-Osman EJ, Veniaminova NA and Merchant JL. Menin and JunD regulate gastrin gene expression through proximal DNA elements. Am J Physiol Gastrointest Liver Physiol. 2011; 301: G783–790.

6. Dreijerink KMA, Ozyerli-Goknar E, Koidl S et al. Multi-omics analyses of MEN1 missense mutations identify disruption of menin-MLL and menin-JunD interactions as critical requirements for molecular pathogenicity. Epigenetics Chromatin. 2022; 15: 29.

7. Anlauf M, Perren A, Henopp T et al. Allelic deletion of the MEN1 gene in duodenal gastrin and somatostatin cell neoplasms and their precursor lesions. Gut. 2007; 56: 637–644.

8. Norton JA, Foster DS, Ito T et al. Gastrinomas: Medical or Surgical Treatment. Endocrinol Metab Clin North Am. 2018; 47: 577–601.

9. Jensen RT, Berna MJ, Bingham DB et al. Inherited pancreatic endocrine tumor syndromes: advances in molecular pathogenesis, diagnosis, management, and controversies. Cancer. 2008; 113: 1807–1843.

10. Rosentraeger MJ, Garbrecht N, Anlauf M et al. Syndromic versus non-syndromic sporadic gastrin-producing neuroendocrine tumors of the duodenum: comparison of pathological features and biological behavior. Virchows Arch. 2016; 468: 277–287.

11. Kimura N, Hirata Y, Iwashiro N, et al. Multiple endocrine neoplasia type 1 with Zollinger-Ellison syndrome: clinicopathological analysis of a Japanese family with focus on menin immunohistochemistry. Front Endocrinol (Lausanne). 2023; 14: 1221514.

12. Anlauf M, Garbrecht N, Henopp T et al. Sporadic versus hereditary gastrinomas of the duodenum and pancreas: distinct clinico-pathological and epidemiological features. World J Gastroenterol. 2006; 12: 5440–5446.

13. Rico K, Duan S, Pandey RL et al. Genome analysis identifies differences in the transcriptional targets of duodenal versus pancreatic neuroendocrine tumours. BMJ Open Gastroenterol. 2021; 8.

14. Wright NA, Pike C and Elia G. Induction of a novel epidermal growth factor-secreting cell lineage by mucosal ulceration in human gastrointestinal stem cells. Nature. 1990; 343: 82–85.

15. Poulsen SS, Nexø E, Olsen PS et al. Immunohistochemical localization of epidermal growth factor in rat and man. Histochemistry. 1986; 85: 389–394.

16. Wee P and Wang Z. Epidermal Growth Factor Receptor Cell Proliferation Signaling Pathways. Cancers (Basel). 2017; 9.

17. Shen T and Guo Q. EGFR signaling pathway occupies an important position in cancer-related downstream signaling pathways of Pyk2. Cell Biol Int. 2020; 44: 2–13.

18. Cao Z, Liao Q, Su M et al. AKT and ERK dual inhibitors: The way forward? Cancer Letters. 2019; 459: 30–40.

19. Evers BM, Rady PL, Sandoval K et al. Gastrinomas demonstrate amplification of the HER-2/neu proto-oncogene. Ann Surg. 1994; 219: 596–601; discussion 602–594.

20. Goebel SU, Iwamoto M, Raffeld M et al. Her-2/neu expression and gene amplification in gastrinomas: correlations with tumor biology, growth, and aggressiveness. Cancer Res. 2002; 62: 3702–3710.

21. Ford MG, Valle JD, Soroka CJ et al. EGF receptor activation stimulates endogenous gastrin gene expression in canine G cells and human gastric cell cultures. J Clin Invest. 1997; 99: 2762–2771.

22. Selvaraj N, Budka JA, Ferris MW et al. Extracellular signal-regulated kinase signaling regulates the opposing roles of JUN family transcription factors at ETS/AP-1 sites and in cell migration. Mol Cell Biol. 2015; 35: 88–100.

23. La P, Desmond A, Hou Z et al. Tumor suppressor menin: the essential role of nuclear localization signal domains in coordinating gene expression. Oncogene. 2006; 25: 3537–3546.

24. Xing B, Ma J, Jiang Z et al. GLP-1 signaling suppresses menin’s transcriptional block by phosphorylation in β cells. J Cell Biol. 2019; 218: 855–870.

25. Duan S, Sheriff S, Elvis-Offiah UB et al. Clinically Defined Mutations in MEN1 Alter Its Tumor-suppressive Function Through Increased Menin Turnover. Cancer Res Commun. 2023; 3: 1318–1334.

26. Schmittgen TD and Livak KJ. Analyzing real-time PCR data by the comparative C(T) method. Nat Protoc. 2008; 3: 1101–1108.

27. La P, Schnepp RW, C DP et al. Tumor suppressor menin regulates expression of insulin-like growth factor binding protein 2. Endocrinology. 2004; 145: 3443–3450.

28. Zeng Z, Zhang Q, Liang T et al. Hsp70 incompletely disaggregates misfolded K488X-menin to promote tumourigenesis in a family with multiple endocrine neoplasia type 1. Cell Signal. 2025; 130: 111681.

29. Cao Y, Liu R, Jiang X et al. Nuclear-cytoplasmic shuttling of menin regulates nuclear translocation of {beta}-catenin. Mol Cell Biol. 2009; 29: 5477–5487.

30. Sundaresan S, Meininger CA, Kang AJ et al. Gastrin Induces Nuclear Export and Proteasome Degradation of Menin in Enteric Glial Cells. Gastroenterology. 2017; 153: 1555–1567.e1515.

31. Oliveira TF, Ferreira HB, Lima LHD et al. A Novel Mutation in a Family With Multiple Endocrine Neoplasia Type 1 and Aggressive Pancreatic Neuroendocrine Tumors. AACE Clin Case Rep. 2025; 11: 126–130.

32. Shiotani A and Merchant JL. cAMP regulates gastrin gene expression. Am J Physiol. 1995; 269: G458–464.

33. Singh B, Carpenter G and Coffey RJ. EGF receptor ligands: recent advances. F1000Res. 2016; 5.

34. Kidd M, Schimmack S, Lawrence B et al. EGFR/TGFα and TGFβ/CTGF Signaling in Neuroendocrine Neoplasia: Theoretical Therapeutic Targets. Neuroendocrinology. 2013; 97: 35–44.

35. Corbo V, Dalai I, Scardoni M et al. MEN1 in pancreatic endocrine tumors: analysis of gene and protein status in 169 sporadic neoplasms reveals alterations in the vast majority of cases. Endocr Relat Cancer. 2010; 17: 771–783.

36. Shah T, Hochhauser D, Frow R et al. Epidermal Growth Factor Receptor Expression and Activation in Neuroendocrine Tumours. Journal of Neuroendocrinology. 2006; 18: 355–360.

37. Di Florio A, Sancho V, Moreno P et al. Gastrointestinal hormones stimulate growth of Foregut Neuroendocrine Tumors by transactivating the EGF receptor. Biochim Biophys Acta. 2013; 1833: 573–582.

38. Duan S, Twer AB, Zinkeng A et al. Hedgehog signaling drives glial cell plasticity and oncogenic reprogramming in gastroenteropancreatic neuroendocrine neoplasms. Molecular Cancer. 2026; 25: 85.

39. Francis J, Lin W, Rozenblatt-Rosen O et al. The menin tumor suppressor protein is phosphorylated in response to DNA damage. PLoS One. 2011; 6: e16119.

40. Irvine F, Pyne NJ and Houslay MD. The phorbol ester TPA inhibits cyclic AMP phosphodiesterase activity in intact hepatocytes. FEBS Lett. 1986; 208: 455–459.

41. Vuorenpää A, Ammendrup-Johnsen I, Jørgensen TN et al. A kinome wide screen identifies novel kinases involved in regulation of monoamine transporter function. Neurochem Int. 2016; 98: 103–114.

42. Zimmermann B, Chiorini JA, Ma Y et al. PrKX is a novel catalytic subunit of the cAMP-dependent protein kinase regulated by the regulatory subunit type I. J Biol Chem. 1999; 274: 5370–5378.

43. Gonzalez FA, Raden DL and Davis RJ. Identification of substrate recognition determinants for human ERK1 and ERK2 protein kinases. J Biol Chem. 1991; 266: 22159–22163.

44. McKenna M, Balasuriya N, Zhong S et al. Phospho-Form Specific Substrates of Protein Kinase B (AKT1). Front Bioeng Biotechnol. 2020; 8: 619252.

45. Wang W, Johansson HE, Bergholm UI et al. Expression of c-Myc, TGF-alpha and EGF-receptor in sporadic medullary thyroid carcinoma. Acta Oncol. 1997; 36: 407–411.

46. Ezzat S. The role of hormones, growth factors and their receptors in pituitary tumorigenesis. Brain Pathol. 2001; 11: 356–370.

47. Höpfner M, Schuppan D and Scherübl H. Treatment of gastrointestinal neuroendocrine tumors with inhibitors of growth factor receptors and their signaling pathways: recent advances and future perspectives. World J Gastroenterol. 2008; 14: 2461–2473.

48. Doleschal B, Petzer A and Rumpold H. Current concepts of anti-EGFR targeting in metastatic colorectal cancer. Front Oncol. 2022; 12: 1048166.

49. Kasi PM, Afable MG, Herting C et al. Anti-EGFR Antibodies in the Management of Advanced Colorectal Cancer. The Oncologist. 2023; 28: 1034–1048.

50. Xiao Z, Xu H, Strosberg JR et al. EGFR is a potential therapeutic target for highly glycosylated and aggressive pancreatic neuroendocrine neoplasms. Int J Cancer. 2023; 153: 164–172.

51. Capurso G, Fazio N, Festa S et al. Molecular target therapy for gastroenteropancreatic endocrine tumours: biological rationale and clinical perspectives. Crit Rev Oncol Hematol. 2009; 72: 110–124.

52. Hobday TJ, Mahoney M, Erlichman C et al. Preliminary results of a phase II trial of gefitinib in progressive metastatic neuroendocrine tumors (NET): A Phase II Consortium (P2C) study. Journal of Clinical Oncology. 2005; 23: 4083–4083.

53. Norden AD, Raizer JJ, Abrey LE et al. Phase II trials of erlotinib or gefitinib in patients with recurrent meningioma. J Neurooncol. 2010; 96: 211–217.

54. Yao JC, Fazio N, Singh S et al. Everolimus for the treatment of advanced, non-functional neuroendocrine tumours of the lung or gastrointestinal tract (RADIANT-4): a randomised, placebo-controlled, phase 3 study. Lancet. 2016; 387: 968–977.

55. Anlauf M, Perren A, Meyer CL et al. Precursor lesions in patients with multiple endocrine neoplasia type 1-associated duodenal gastrinomas. Gastroenterology. 2005; 128: 1187–1198.

56. Kim H, Lee JE, Cho EJ et al. Menin, a tumor suppressor, represses JunD-mediated transcriptional activity by association with an mSin3A-histone deacetylase complex. Cancer Res. 2003; 63: 6135–6139.

57. Hughes CM, Rozenblatt-Rosen O, Milne TA et al. Menin associates with a trithorax family histone methyltransferase complex and with the hoxc8 locus. Mol Cell. 2004; 13: 587–597.

